# ECM dimensionality tunes actin tension to modulate the endoplasmic reticulum and spheroid phenotype

**DOI:** 10.1101/2021.07.14.452329

**Authors:** FuiBoon Kai, Guanqing Ou, Richard W. Tourdot, Connor Stashko, Guido Gaietta, Mark F. Swift, Niels Volkmann, Alexandra F. Long, Yulong Han, Hector H. Huang, Jason J. Northey, Andrew M. Leidal, Virgile Viasnoff, David M. Bryant, Wei Guo, Arun P. Wiita, Ming Guo, Sophie Dumont, Dorit Hanein, Ravi Radhakrishnan, Valerie M. Weaver

## Abstract

Primary tissue organoids and cell spheroids recapitulate tissue physiology with remarkable fidelity. We investigated how engagement with a three dimensional laminin-rich extracellular matrix supports the polarized, stress resilient spheroid phenotype of mammary epithelial cells. Cells within a three dimensional laminin-rich extracellular matrix decreased and redistributed the actin crosslinker filamin to reduce their cortical actin tension. Cells with low cortical actin tension had increased plasma membrane protrusions that promoted negative plasma membrane curvature and fostered protein associations with the plasma membrane, consistent with efficient protein secretion. By contrast, cells engaging a laminin-rich extracellular matrix in two dimensions had high filamin-dependent cortical actin tension, exhibited compromised endoplasmic reticulum function including increased expression of PKR-like Endoplasmic Reticulum Kinase signaling effectors, and had compromised protein secretion. Cells with low filamin-mediated cortical actin tension and reduced endoplasmic reticulum stress response signaling secreted, and assembled, a polarized endogenous basement membrane and survived better, and their spheroids were more resistant to exogenous stress. The findings implicate filamin-dependent cortical actin tension in endoplasmic reticulum function and highlight a role for mechanics in organoid homeostasis.

## Introduction

Tissue fragments cultured within a three dimensional (3D) reconstituted basement membrane (rBM) as organoids retain many of the differentiated features of their native tissue including their apical-basal polarity, vectorial protein secretion, and resistance to exogenous stress^1–3^. Recapitulation of the tissue-like differentiated spheroid phenotype of primary and immortalized cells, and the retention of their long-term viability and stress resilience, is similarly supported by culturing the cells within a 3D rBM^2, 4^. Nevertheless, it remains unclear how a 3D rBM directs this tissue-specific structure and function.

Primary and immortalized mammary epithelial cells (MECs) embedded within a 3D rBM assemble into growth-arrested, apoptosis-resistant spheroids, secrete and assemble an endogenous basement membrane and express tissue-specific differentiated gene expression^4^. The tissue-specific differentiation of MECs depends upon the engagement of laminin receptors and a rounded cell phenotype that direct the signaling required to induce differentiated gene expression^5^. A compliant extracellular matrix (ECM), in turn, supports MEC rounding and stabilizes the cell-cell adhesions required to build and maintain the differentiated spheroid structure^6, 7^. Importantly however, a two-dimensional (2D) compliant laminin-rich rBM fails to support the establishment of a stress resilient, polarized 3D spheroid surrounded by an endogenous basement membrane. Why secretion and deposition of a polarized basement membrane and acquisition of long-term viability and treatment resistance in MEC spheroids requires interaction with a compliant 3D laminin-rich rBM is poorly understood.

Organelles are subcellular structures that execute specialized cellular functions such as energy production, and protein secretion, recycling and degradation required for cell and tissue homeostasis. The endoplasmic reticulum (ER) is a multifunctional organelle essential for protein folding that is critical for secretion and for the stress regulation required to support normal cell and tissue function. The ER is particularly critical for calcium storage, synthesis and folding of nascent transmembrane and secretory proteins, and for lipid metabolism. ER function is compromised when ER calcium levels are depleted and this results in protein misfolding, induction of ER stress and activation of ER stress response^8^. Not surprisingly, cells have developed a sophisticated surveillance system to monitor and ensure efficient ER activity. Upon calcium depletion in the ER, the ER quickly establishes ER-plasma membrane contact sites that activate store-operated calcium entry (SOCE) to replenish intracellular calcium storage to restore ER function^9^. Thus, the ER plays a critical role in maintaining cell and tissue homeostasis and operate key mechanisms to facilitate this function. Given the profound impact of the ER on protein secretion and cell stress regulation, we asked whether ECM dimensionality, per se, could regulate endogenous basement membrane deposition and direct stress resilience in 3D spheroids by modulating ER structure and/or function and if so how?

## Results

### A 3D laminin/rBM increases expression of molecules implicated in regulating endoplasmic reticulum function

In marked contrast to MECs grown as 2D monolayers on top of a thin layer of a laminin-rich rBM, immortalized nonmalignant MCF10A MECs grown within a 3D laminin-rich rBM generate growth-arrested spheroids^10^. Growth-arrested polarized MEC spheroids within a 3D laminin-rich rBM increase the expression of genes implicated in collagen-containing ECM proteins and plasma membrane surface protein localization (Fig. 1a and b)^4, 11^. The spheroids generated within a 3D laminin-rich rBM also acquire apical-basal polarity (as indicated by cell-cell localized E-cadherin, apical-lateral ZO-1 and basal-lateral β4 integrin) and demonstrate vectorial secretion of proteins (Fig. 1c; top panels). This includes the basal deposition of an endogenous basement membrane, as indicated by type IV collagen and laminin-5, and the apical secretion of whey acidic protein and β-casein (Fig. 1c; bottom panels)^1, 4, 12^. Furthermore, the spheroids generated within a laminin-rich rBM are highly resistant to exogenous stresses including treatment with agents such as Paclitaxol, Trail, Doxorubicin, as well as gamma radiation exposure as compared to MECs interacting with a 2D laminin-rich rBM (Fig 1d). Indeed, we found that MECs interacting with 2D laminin-rich rBM exhibit an ER stress phenotype when treated with agents such as Trail, as illustrated by enhanced expression of genes reflecting the ER chaperone complex, response to ER stress and PERK-mediated UPR (Fig. 1e).

**Figure 1.**
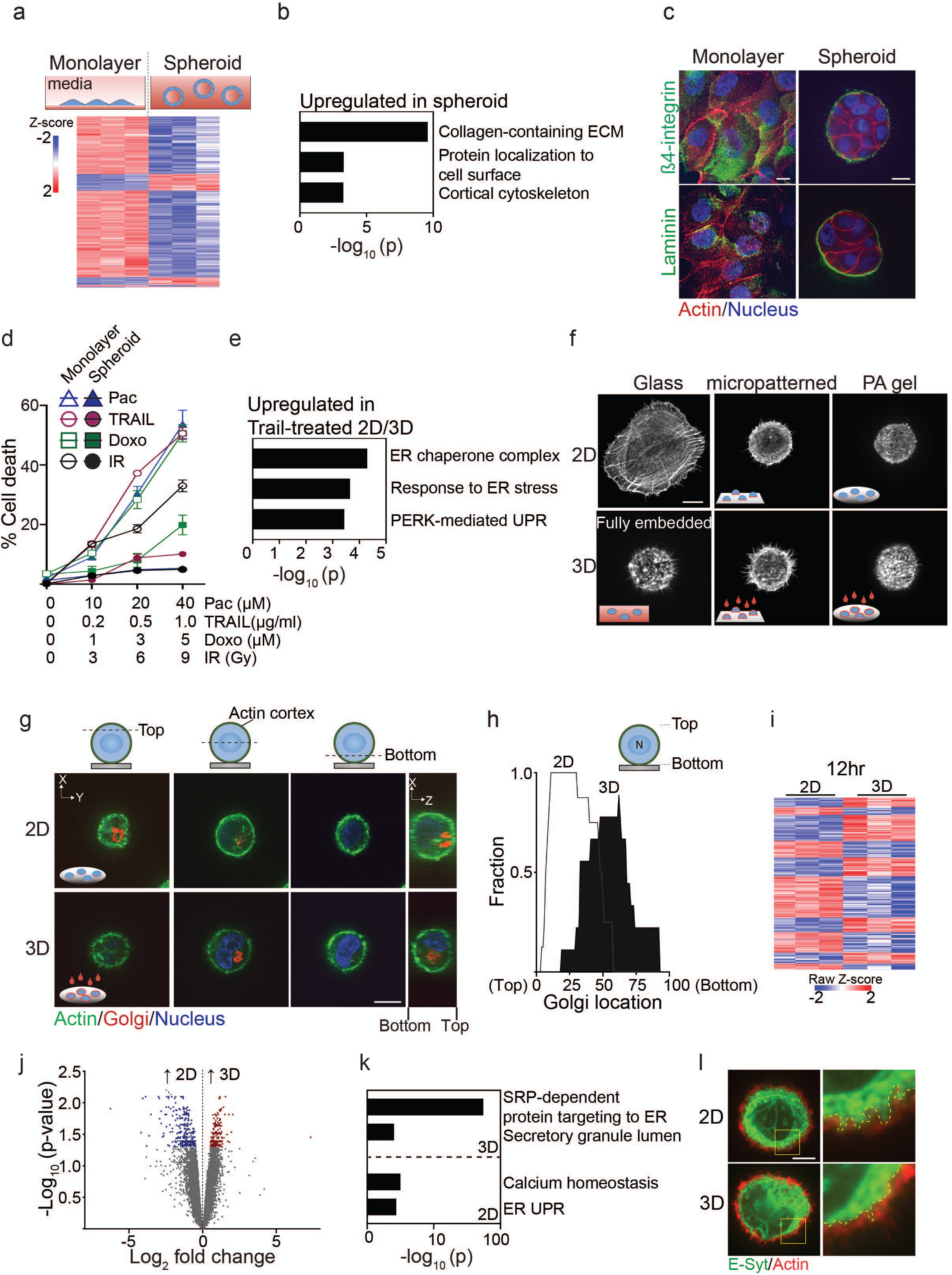
A 3D laminin/rBM increases expression of molecules implicated in regulating ER function. **(a)** (Top panel) Representative immunofluorescence microscopy images of MECs grown as monolayers on a rigid rBM (two-dimensional culture, 2D; left) or within a compliant rBM (three-dimensional culture, 3D) to generate multicellular spheroid structures with apical-basal polarity (right). Scale bar, 10 μm. (Bottom panel) Heatmap of microarray analysis of gene expression in MECS cultured either as a 2D monolayer or as spheroids. Expression of the top 500 differentially expressed genes between the two conditions are shown (n=3). **(b)** Gene Ontology (GO) analysis of genes upregulated in 3D spheroids. (**c)** Samples were stained with antibodies for laminin or β4-integrin (green). The actin and nucleus were counterstained with phalloidin (red) and DAPI (blue), respectively. Scale bar, 10 μm. **(d)** MECs plated as monolayers on a rigid rBM (2D) or as spheroids within rBM (3D) were treated with increasing doses of Paclitaxol (Pac), TRAIL, Doxorubicin (Doxo) and irradiation (IR). Percent cell death was quantified by immunofluorescence as percentage of cells staining positively for cleaved caspase-3 48 hr post-treatment. (Mean ± s.e.m.; n=3). **(e)** GO analysis of genes significantly upregulated in TRAIL-treated MEC monolayers (2D) relative to TRAIL-treated spheroid cultures (3D). **(f)** Representative immunofluorescence microscopy images of MECs plated as single cells on rigid glass coverslips (glass) or fully embedded within rBM and stained with phalloidin to reveal F-actin organization. Cell spreading was inhibited by plating cells on either laminin-111 conjugated, 10-μm micropatterned glass (2D/micropatterned, rigid substrate) or on compliant 75 Pa rBM-laminated polyacrylamide (PA) gels (2D/PA, soft substrate). The single 2D cells were overlaid with either purified laminin-111 or rBM to create a 3D ECM microenvironment. Images show maximum intensity z-projections of confocal stacks for F-actin phalloidin staining. Scale bar, 10 μm. **(g)** Representative immunofluorescence microscopy images of MECs stably expressing recombinant mCherry-tagged golgi marker (GalT; red) ligated with rBM in either 2D or 3D. The actin cortex and nucleus were counterstained with phalloidin (green) and DAPI (blue), respectively. Images show the cross-sectional view of each cell compartment (dashed lines; xy plane) and side view of confocal stacks (xz plane) in the individual MECs. **(h)** Golgi staining was assessed within nonspread MECs ligated with rBM in 2D and 3D and values were plotted as a function of subcellular localization. (2D, n=8; 3D, n=9) **(i)** Heatmap of RNA-seq experiment from MECs ligated with rBM in 2D and 3D 12 hours post-plating. Data show the expression of the top 489 genes that are differentially expressed between the 2D and 3D rBM conditions. (n=3). **(j)** Volcano plot of differentially expressed genes from RNA-seq of MECs ligated to rBM in 2D and 3D harvested 12 hours post-plating. Significantly downregulated genes (blue; log 2> 0.5) and upregulated genes (red; log 2> 0.5) are highlighted. **(k)** GO analysis showed functional enrichment of genes significantly upregulated in nonspread MECs ligated to rBM in 3D relative to those interacting with a rBM in 2D. **(l)** Representative immunofluorescence microscopy images of MECs ligated to 2D or 3D rBM and stained with an antibody targeting Extended-synaptotagmin (E-Syt), an ER-anchored protein that mediates the tethering of the ER to the plasma membrane. F-actin was counterstained using phalloidin (red). Color images show whole cells and boxed areas are magnified (bottom right; 2D and 3D panels). The ER border is marked with a yellow dashed line. Scale bar, 10 μm.

To clarify whether ECM ligation dimensionality modulates spheroid phenotype by altering ER function we studied the impact of engaging MECs with a laminin-rich rBM in 2D versus 3D. To avoid potential contributions induced by cell-cell junctions and multicellularity on ER function we applied a single cell assay in which laminin ligation, and cell spreading were restricted and ECM ligation in 2D versus 3D was controlled. We maintained cell rounding, which is a feature of a MEC organoid engaging a 3D laminin-rich rBM, by limiting cell spreading by plating single cells on top of either micropatterned (10 μm) laminin-conjugated borosilicate glass or on a compliant (75 Pascal) laminin-rich rBM-laminated polyacrylamide (PA) gel (Fig. 1f; compare 2D nonpatterned glass to micropatterned glass and PA gels). The third ECM dimension was induced by overlaying the non-spread MECs with either purified laminin-111 or a laminin-rich rBM (Fig. 1f; compare 2D to 3D). We evaluated the impact of ECM compliance on cell phenotype by comparing actin organization in the non-spread MECs plated on the rigid micropatterned glass to the cells that were plated on top of the compliant PA gels and that were interacting with the laminin-rich rBM in 2D versus those interacting with the laminin-rich rBM in 3D. To begin with, we noted that all of the rounded single cells reconstituted an actin phenotype highly reminiscent to that demonstrated by single MECs fully embedded within a 3D laminin-rich rBM, whether they were plated on top of a compliant laminin-rich rBM or on top of a micropatterned laminin-conjugated rigid glass substrate (Fig. 1f). The impact of ECM dimensionality was verified by showing loss of forced apical-basal polarity in the MECs plated either on the laminin-rich rBM PA gels overlaid with either purified laminin or a laminin-rich rBM. The non-polarized phenotype was illustrated by the absence of Golgi apparatus orientation in the MECs ligated with the laminin-rich rBM in 3D and the uniform apical localization of this organelle in the MECs plated on top of the 2D laminin-rich rBM (Fig. 1g & h).

Global transcriptome analysis revealed that within 12 hours, out of 489 differentially expressed genes, 201 genes were induced, and 288 genes were repressed in the single non-spread MECs interacting with the laminin-rich rBM in 3D, as compared to the single non-spread MECs interacting with the laminin-rich rBM in 2D (Fig. 1i & j). Gene set enrichment analysis revealed that the single non-spread MECs with the cortically organized actin that engaged the laminin-rich rBM in 3D were enriched for genes implicated in regulation of protein insertion into the ER. By contrast, the MECs encountering the 2D laminin-rich rBM were enriched for genes implicated in the unfolded protein response, which is a cellular stress response activated by ER stress. (Fig. 1k). Examination of ER morphology confirmed effects on ER function by revealing an ER network that appears to penetrate through the actin cortex when the MECs interact with the laminin-rich rBM in 2D that was less obvious in the MECs engaging the laminin-rich rBM in 3D (Fig. 1l). The findings suggest that how a cell ligates its ECM may regulate its ER organization and function.

### Ligation of a laminin-rich rBM in 3D alters ER function

We next explored whether cell ligation to its ECM in two versus three dimensions influences its ER organization and function. Building upon our GO analysis (Fig. 1k), we examined the impact of ligating the laminin-rich rBM in 2D versus 3D on ER function by monitoring effects on secretory protein trafficking. To begin with, RT-PCR analysis confirmed that the MECs interacting with the laminin-rich rBM in 3D had much higher levels of SEC61 expression, which is a subunit of the channel-forming translocon complex responsible for targeting proteins to the secretory pathway by directing their insertion either into the ER membrane or lumen (Fig. 2a). By expressing an EGFP-tagged temperature-sensitive vesicular stomatitis virus G protein (VSVG-ts045) in MECs, we then tested whether those MECs ligating the laminin-rich rBM in 2D versus 3D showed differences in secretory protein trafficking. Temperature induced pulse chase monitoring for secretory protein trafficking revealed a consistent and significant increase in plasma membrane-associated VSVG-ts045 in the MECs interacting with the laminin-rich rBM in 3D consistent with enhanced secretory protein trafficking (Fig. 2b & 2c).

**Figure 2.**
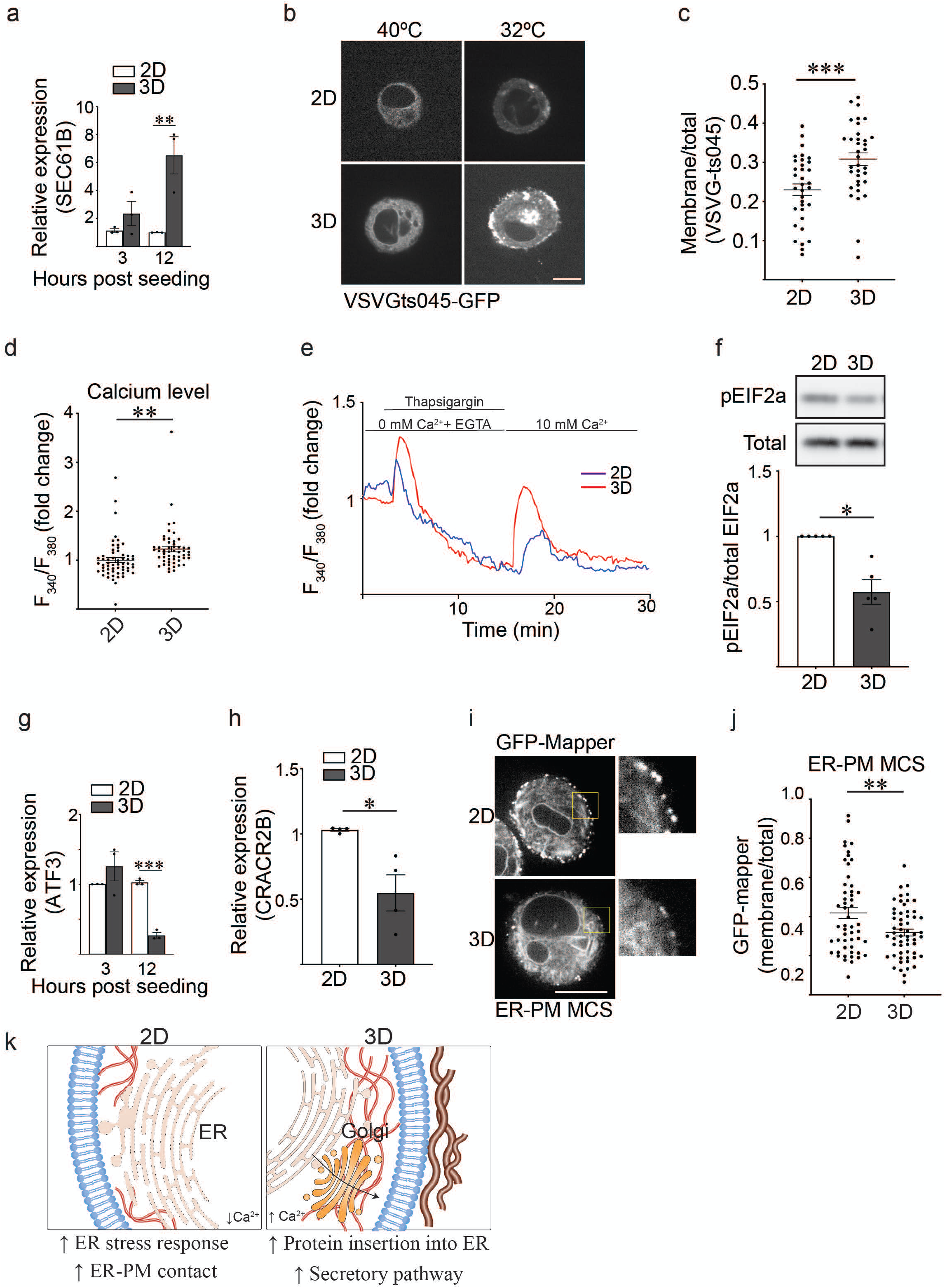
Ligation of a laminin-rich rBM in 3D alters ER function. **(a)** Bar graphs showing qPCR quantification of SEC61B (Mean ± s.e.m.; n=3) level in MECs ligated with rBM in 2D and 3D 12 hours post-plating. **(b)** MECs expressing VSVGts045-EGFP were ligated to rBM in 2D or 3D and incubated at 40°C for 16 hr. Cells were shifted to 32°C for 2 hr to release VSVGts045-EGFP into the secretory pathway. Shown are representative fluorescence microscopy images of MECs expressing VSVGts045-EGFP 2 hr post incubation at 32°C. Scale bar, 10 μm. **(c)** Scatter plot showing secretory trafficking efficiency quantified based on the fluorescence intensity of VSVGts045 at the plasma membrane/total VSVGts045 fluorescence intensity in the cells. (Mean ± s.e.m.; n=34 cells). **(d)** Ratiometric calcium indicator Fura-2 measurements showing basal [Ca^2+^] levels in MECs ligated to a rBM in 2D or 3D (Mean ± s.e.m.; 2D, n=58; 3D, n=53). **(e)** Line graph showing the changes in [Ca^2+^] levels in MECs under different treatment. Fura-2-loaded MECs ligated to 2D or 3D rBM were incubated with Ca^2+^-free HBSS buffer, challenged with 2 uM Thapsigargin, and replenished with 10 mM Ca^2+^ at the indicated time. (Mean; 2D, n=7; 3D, n=6). **(f)** Graphs showing quantification of the level of phosphorylated EIF2a (pEIF2a) and total EIF2a in MECs ligated to a rBM in 2D or 3D as assessed via immunoblot (Mean ± s.e.m.; n = 5). **(g)** Bar graphs showing qPCR of the relative level of ATF3 mRNA in MECs ligated with rBM in 2D and 3D (Mean ± s.e.m.; n = 3). **(h)** Bar graphs showing qPCR of the relative level of CRACR2B mRNA in MECs ligated with rBM in 2D and 3D (Mean ± s.e.m.; n = 4). **(i)** Representative fluorescence microscopy images of GFP-MAPPER (a reporter of ER-PM junctions) in MECs ligated with rBM in 2D or 3D. Scale bar, 10 μm. **(j)** Quantification of the amount of GFP-MAPPER in MECs ligated with rBM in 2D or 3D. The abundance of ER-PM contact sites in MECs was quantified as plasma membrane fluorescence intensity relative to total fluorescence intensity (Mean ± s.e.m.; n=52; 3D, n=57). **(k)** Graphical schematic showing how ECM dimensionality can influence ER structure/function.

Our GO analysis indicated that MECs ligating a laminin-rich rBM in 2D upregulated genes implicated in regulating calcium homeostasis and ER stress response. Thus, we next investigated whether the nature of laminin-rich rBM ligation by the MECs differentially regulated intracellular calcium homeostasis. FURA-2 analysis revealed that the MECs ligating the laminin-rich rBM in 2D not only had lower resting intracellular calcium content (Fig. 2d) but also exhibited compromised calcium regulation, as shown by a truncated amplitude of intracellular calcium release following thapsigargin treatment, and store-operated calcium entry (SOCE) (Fig. 2e). Loss of calcium homeostasis can lead to accumulation of unfolded protein in the ER and can activate a ER stress response. We therefore focused on PERK-pEIF2a-ATF4-ATF3 signaling axis because ATF3 is one of the top candidate genes from our RNA-seq we found to be significantly upregulated in the single MECs ligating the laminin-rich rBM in 2D. Consistently, the MECs interacting with the laminin-rich rBM in 2D had higher levels of phosphorylated EIF2a (pEIF2a; Fig. 2f) and the stress regulator ATF3 (Fig. 2g) when compared to the MECs interacting with the laminin-rich rBM in 3D.

In response to calcium depletion, cells activate SOCE at the ER-PM contact sites to replenish their intracellular calcium stores. Our RNA-seq analysis revealed that the MECs interacting with a laminin-rich rBM in 2D as compared to those interacting with the same rBM in 3D, expressed high levels of CRACR2B, which regulates Ca^2+^ release-activated Ca^2+^ channels that mediate SOCE (validated in Fig. 2h). We explored this phenotype further by examining the presence of ER-PM contact sites in the MECs interacting with the laminin-rich rBM in 2D and 3D. Using the GFP-MAPPER marker which reports on ER-PM junctions, we observed that the MECs interacting with the laminin-rich rBM in 2D not only had compromised calcium regulation but also exhibited a significant increase in the number of ER-PM contact sites (Fig. 2i & 2j). These data confirm that how a cell ligates its ECM does influence ER structure and function (Fig. 2k).

### Ligation of a laminin-rich rBM in 3D modulates filamin to alter ER function

ER-PM contact site assembly is mediated by interaction between the actin cross-linker filamin and the ER stress sensor PERK^13^. We therefore assessed whether ligating the ECM in 2D versus 3D influenced ER organization and function by modulating filamin levels and cellular distribution. Immunostaining revealed prominent apically localized filamin aggregates in the MECs engaging the laminin-rich rBM in 2D, whereas by contrast filamin was diffusely localized throughout the MECs interacting with the laminin-rich rBM in 3D (Fig. 3a). Immunoblot analysis further revealed that filamin protein levels were significantly lower in the MECs interacting with the ECM in 3D (Fig. 3b). Moreover and consistent with a functional link between filamin and ER-PM junctions, shRNA-mediated knockdown of filamin reduced the number of ER-PM junctions in the MECs interacting with the laminin-rich rBM in 2D as indicated by the GFP-mapper reporter (Fig. 3c & d). Consistent with a functional link between filamin-mediated ER-PM junctions and ER function, depleting filamin in the MECs ligating the laminin-rich rBM in 2D ameliorated ER stress signaling, as indicated by lower levels of phosphorylated EIF2a protein and reduced expression of the stress regulator ATF3 in the MECs (Fig 3e & f). Conversely, overexpression of filamin in the MECs interacting with the laminin-rich rBM increased their level of pEIF2a implying an induction of ER-mediated cell stress (Fig. 3g). The findings suggest that a cell modulates its phenotype in response to engagement with an ECM in either 2D or 3D by modulating filamin-ER interactions to regulate ER organization and function (Fig. 3h).

**Figure 3.**
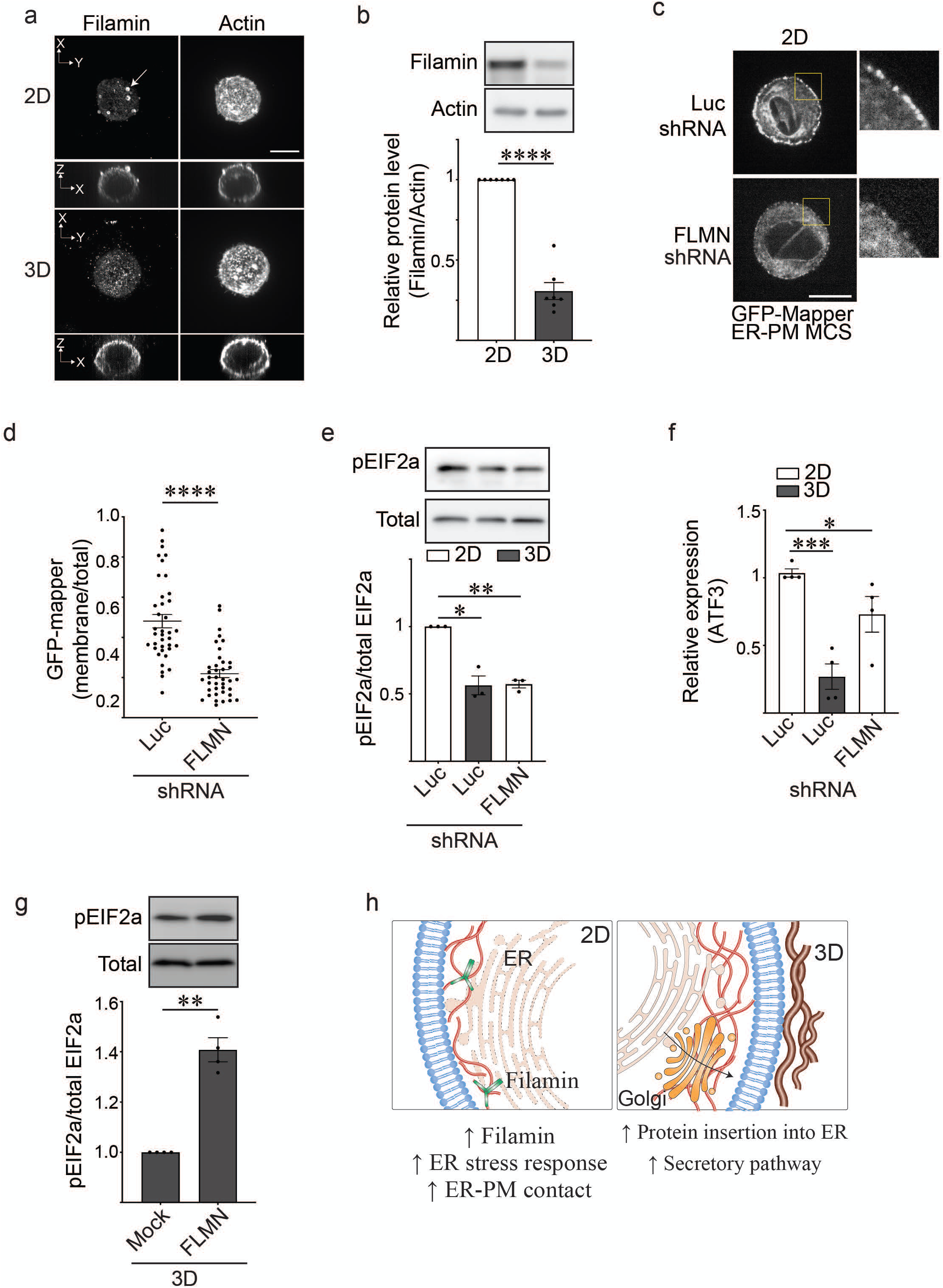
Ligation of a laminin-rich rBM in 3D modulates filamin to alter ER function. **(a)** Representative immunofluorescence microscopy images of MECs ligated with either a rBM in 2D or 3D and stained with an antibody targeting filamin. F-actin was counterstained with phalloidin. Images show maximum intensity z-projections of xy confocal stacks and side view of xz confocal stacks of MECs. Arrows indicate filamin aggregates in the individual MECs ligated to rBM in 2D. Scale bar, 10 μm. **(b)** The expression level of filamin in MECs ligated to rBM in 2D and 3D was assessed via immunoblot and quantified relative to total cellular actin (Mean ± s.e.m.; n = 7). **(c)** Representative fluorescence microscopy images of GFP-MAPPER (a reporter of ER-PM junctions) in MECs stably expressing an shRNA targeting filamin (FLMN) or luciferase (Luc) ligated to rBM in 2D. Scale bar, 10 μm. **(d)** Quantification of the amount of GFP-MAPPER in MECs stably expressing an shRNA targeting filamin (FLMN) or luciferase (Luc) ligated to rBM in 2D. The abundance of ER-PM contact sites in MECs was quantified as plasma membrane fluorescence intensity relative to total fluorescence intensity. **(e)** Bar graphs showing quantification of the level of pEIF2a and EIF2a in MECs stably expressing an shRNA targeting filamin (FLMN) or luciferase (Luc) ligated to rBM in 2D or 3D as measured by immunoblotting and quantified relative to total cellular protein. (Mean ± s.e.m.; n=3). **(f)** Bar graphs showing qPCR of the relative level of ATF3 mRNA in MECs ligated with rBM in 2D and 3D expressing either an shRNA targeting FLMN or a control luciferase vector (Luc). (Mean ± s.e.m.; n = 4). **(g)** Bar graphs showing quantification of the level of pEIF2a and EIF2a in MECs ligated to rBM in 3D treated with EtOH (mock) or doxycycline (FLMN) to induce filamin expression (Mean ± s.e.m.; n=3). **(h)** Graphical schematic showing how ECM dimensionality can influence ER function via filamin.

### Ligation of a laminin-rich rBM in 3D alters ER function by reducing actin tension

Filamin is a mechanosensitive actin-cross-linker^14^. Given our findings that the dimensionality of ECM ligation by a cell influences the expression and spatial organization of cellular filamin, we asked whether ECM ligation dimensionality influenced actin cytoskeletal tension. We used Atomic Force Microscopy (AFM) to measure the tension of the actin cortex in non-spread MECs plated on top of laminin-111 coated micro-patterned borosilicate glass (10μm), and generated the third dimension of ECM binding using a dilute concentration of laminin-111 to limit non-specific binding and potential measurement interference. AFM force-distance curves confirmed that the laminin-111 overlay did not impair AFM indentation measurements (Fig. S4a). AFM indentation revealed that the MECs engaging laminin in 2D had a significantly stiffer actin cortex than the same MECs overlaid with laminin-111 to generate the third dimension of ECM engagement (Fig. 4a; compare 2D to 3D). Treatment of the MECs with blebbistatin confirmed that actin cortex stiffness was due to myosin-II activity (compare 2D to 3D with and without blebbistatin treatment). Furthermore, inducing ROCK activity to increase actomyosin tension restored cortical actin tension in the MECs engaging the ECM in 3D towards that measured in the MECs engaging the ECM in 2D (compare 2D to 3D + ROCK). Consistent with our earlier observations that increasing cellular filamin prevented the drop in cortical actin stiffness induced in the MECs by engagement with the ECM in 3D (Fig. 4a; compare 3D to 3D + FLMN). Furthermore, Traction Force Microscopy (TFM) measurements, that quantify actomyosin contractility, revealed that MECs ligating the laminin-rich rBM in 3D exerted lower traction stress against the basal ECM substrate, likely through a redistribution of traction forces around the cell cortex (Fig. 4b Fig. S4b). Moreover, inhibiting myosin-II activity by treating the cells with blebbistatin, uniformly abolished traction stresses in the MECs regardless of ECM ligation context (Fig. 4b). To assess if cortical actin tension associated with changes in cellular rheology, we used optical tweezers to measure the cytoplasmic modulus in the MECs ^15, 16^. The measurements revealed that the MECs engaging the laminin-rich rBM in 3D had a lower cytoplasmic modulus as compared to the MECs engaging the laminin-rich rBM in 2D (Fig. 4c).

**Figure 4.**
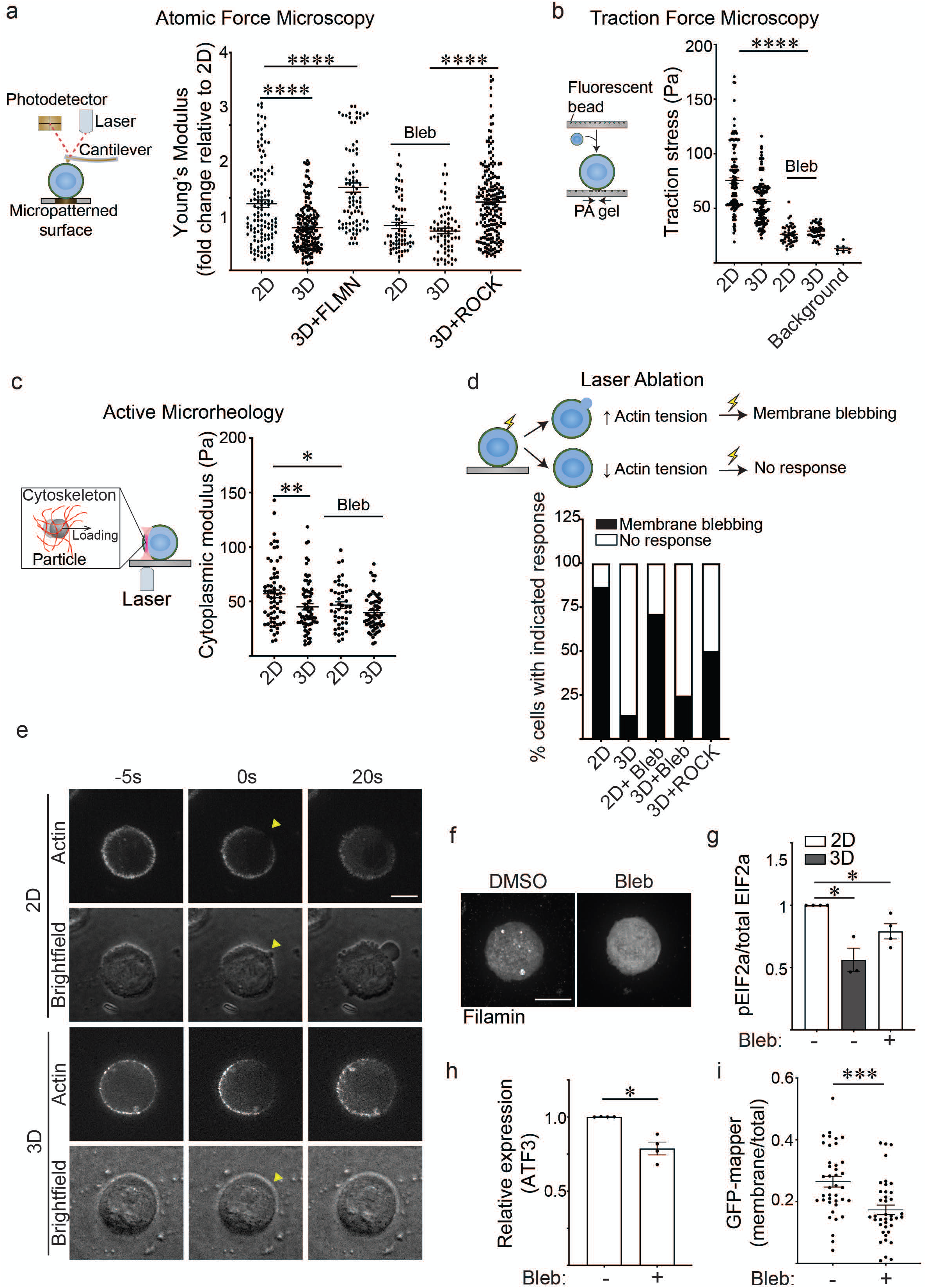
Ligation of a laminin-rich rBM in 3D alters ER function by reducing actomyosin tension. **(a)** (Left) Cartoon depicting the principle of Atomic Force Microscopy (AFM). A cantilever at the end of the microscope probe is deflected when it is in contact with the cell surface. Cell cortex-mediated resistance to indentation alters the path of the laser beam focused on the cantilever that is then reflected onto a photodetector to enable measurement of cellular cortical actin tension. (Right) AFM was used to measure the cortical actin tension for MECs ligated to a laminin-111 substrate either in 2D or 3D. The MECs were treated with doxycycline to induce filamin expression (FLMN), with or without the myosin II inhibitor blebbistatin (Bleb) to reduce cortical actin tension, or the cells were engineered to express an inducible activated ROCK to increase cortical actin tension. The MECs were indented using a 2-µm beaded tip on the AFM cantilever and the Hertz model was used to fit each indentation curve to extract the Young’s modulus of the cell cortex (Mean ± s.e.m from at least n=30 cells from three independent experiments). **(b)** (Left) Cartoon depicting the principle of traction force microscopy. MECs are plated on compliant 400 Pa polyacrylamide gels containing 100 nm fluorescent beads (close to the cell-polyacrylamide gel interface). Traction stresses are calculated based on the bead displacement induced by substrate deformation and relaxation. (Right) Quantification of the traction stresses in individual MECs ligated with rBM in 2D and 3D and treated with or without the myosin II inhibitor blebbistatin (Bleb). (Mean ± s.e.m.; 2D, n= 139; 3D, n=124; 2D+Bleb, n=42; 3D+Bleb, n=43). The background bead displacement in the traction force analysis was measured from gel areas that lacked ligated cells (Mean ± s.e.m.; background, n= 8). **(c)** (Left) Cartoon depicting principle of active microrheology used for the measurements. MECs endocytosed 0.5 μm polystyrene particles, which were trapped and oscillated using laser optical tweezers to measure the cytoplasmic modulus. (Right) The cytoplasmic modulus was measured for MECs ligated to a rBM in 2D or 3D and treated with or without blebbistatin (Bleb) to reduce cortical actin tension. For these measurements, the cytoplasmic modulus was calculated based on the slope in the linear range of the normalized force-displacement curve. (Mean ± s.e.m.; 2D, n= 66; 3D, n=69; 2D+Bleb, n=46; 3D+Bleb, n=60 from three independent experiments). **(d)** (Top) Cartoon depicting strategy used to measure cortical tension using laser ablation. Cells with high cortical tension exhibit plasma membrane blebbing when cortical actin is severed by a pulsed laser, whereas cells with lower cortical tension do not. (Bottom) Bar graphs showing quantification of the response of MECs cultured under the indicated conditions following laser ablation. (Mean; 2D, n= 14; 3D, n=15; 2D+Bleb, n=17; 3D+Bleb, n=14; 3D+ ROCK, n=10 cells). **(e)** Representative fluorescence and brightfield images of bleb formation induced by laser ablation in MECs stably expressing LifeAct-RFP. Arrowhead: the site of laser ablation. Scale bar, 10 μm. **(f)** Representative fluorescence microscopy images of MECs ligated with a rBM in 2D and treated with or without blebbistatin (Bleb) and stained with an antibody targeting filamin. Images show maximum intensity z-projections of xy confocal stacks of MECs. Scale bar, 10 μm. **(g)** Bar graphs showing quantification of the level of pEIF2a and EIF2a in MECs ligated with rBM in 2D and 3D and treated with or without blebbistatin (Bleb) as measured by immunoblotting and quantified relative to total cellular protein (Mean ± s.e.m.; n=3). **(h)** Bar graphs showing qPCR of the relative level of ATF3 mRNA in MECs ligated with rBM in 2D and treated with or without blebbistatin (Bleb) (Mean ± s.e.m.; n = 4). **(i)** Quantification of GFP-MAPPER abundance ligated with rBM in 2D and treated with or without blebbistatin (Bleb). The abundance of ER-PM contact sites in MECs was quantified as plasma membrane fluorescence intensity relative to total fluorescence intensity (Mean ± s.e.m.; n = 39 cells per condition).

Cells with high actin tension can nucleate blebs rapidly in response to local laser ablation of the actin cortex^17^. Thus, we used laser ablation to disrupt the actin cortex and monitored the bleb behavior of the untethered plasma membrane to examine whether the context of ECM ligation influences cortical actin tension. Consistently, laser ablation of cortical actin induced rapid bleb formation, in an actomyosin-dependent manner, in the majority of the MECs interacting with the laminin-rich rBM in 2D (Fig. 4d & e). By contrast, laser ablation failed to elicit blebs in most of the MECs interacting with the laminin-rich rBM in 3D. Importantly, however, expressing a constitutively active ROCK to elevate actomyosin activity restored laser ablation-induced membrane bleb activity in the MECs engaging the ECM in 3D, implicating actomyosin tension in this phenotype and ruling out the possibility that the ECM overlay physically impedes bleb formation (Fig. 4d & e; compare 2D, 3D and 3D+ROCK).

To determine whether the context of ECM ligation influences ER function by tuning cortical actin tension, we treated the MECs engaging the laminin-rich rBM in 2D with blebbistatin and assessed whether this treatment reduced ER stress signaling and altered ER organization. Data revealed that inhibiting actomyosin activity eliminated apically localized filamin aggregates (Fig. 4f), reduced the level of EIF2a protein in the MECs engaging the laminin-rich rBM in 2D (Fig. 4g) and simultaneously decreased levels of the stress-regulated transcription factor ATF3 (Fig. 4h). Reducing actomyosin tension also decreased the number of ER-PM junctions presumably reflecting improved ER calcium homeostasis (Fig. 4i). These findings indicate that the dimensionality context of ECM ligation by cells can influence ER function by tuning cortical actin tension.

### Cortical actin tension modulates plasma membrane topology

ER stress can impede protein secretion^18^. Thus, we asked whether actin tension influenced secretory protein trafficking in the MECs engaging the laminin-rich rBM in 3D. Consistently, inhibiting actomyosin tension in the MECs engaging the ECM in 2D reduced ER stress signaling and concomitantly increased SEC61 expression (Fig. 5a), suggesting low cortical tension could modulate secretory protein trafficking by regulating ER function.

**Figure 5.**
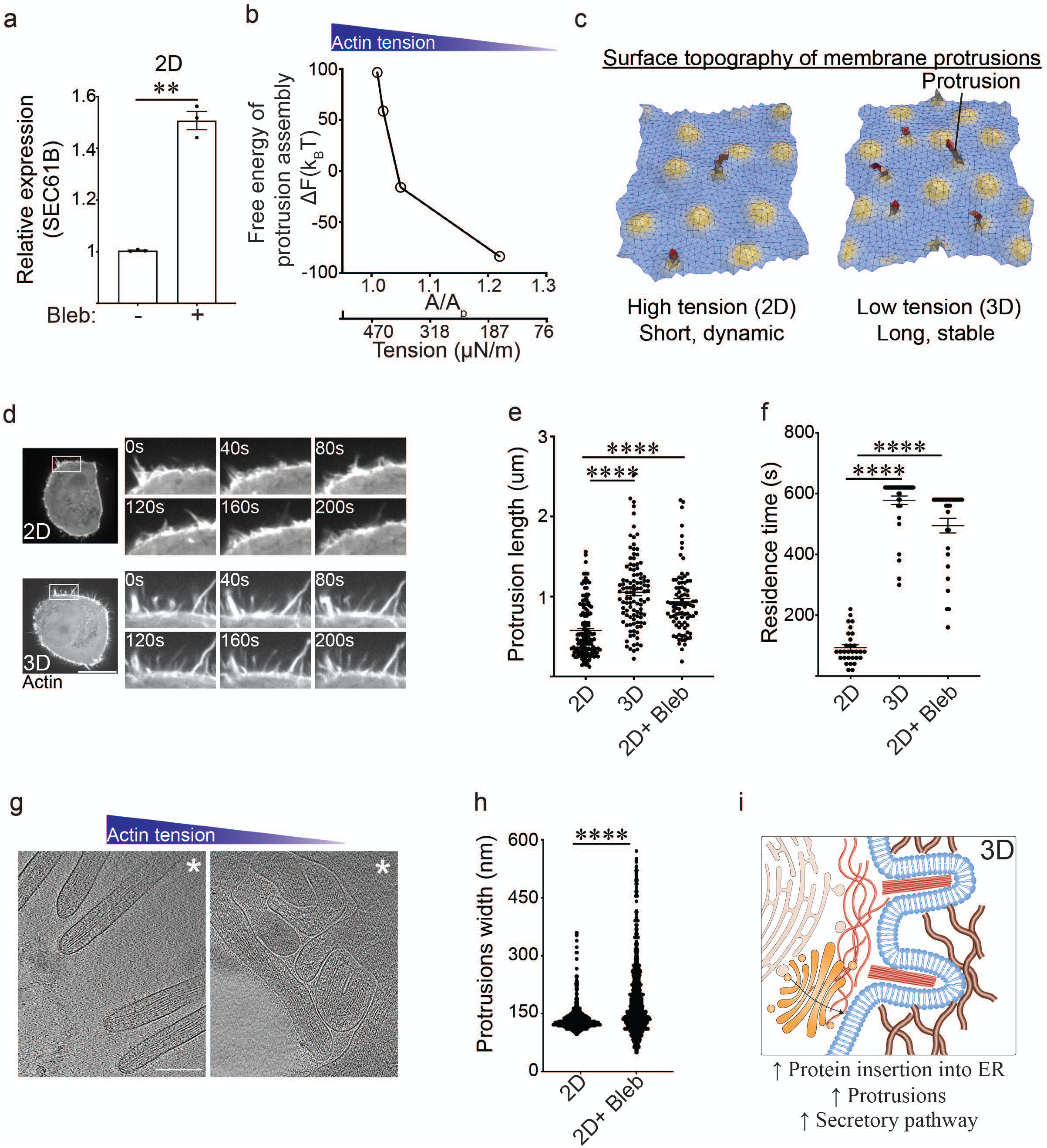
Cortical actin tension modulates plasma membrane topology. **(a)** Bar graphs quantifying SEC61B mRNA levels measured via qPCR in MECs ligated to rBM in 2D with and without blebbistatin (Bleb) treatment (Mean ± s.e.m.; n=3). **(b)** Change in the Helmholtz free energy of membrane protrusion assembly *(ΔF)* as a function of cell membrane excess area (A/A_p_), which is the variable conjugate to the cortical actin tension. **(c)** Representative snapshots from a typical membrane patch with properties of the plasma membrane showing that protrusion length is higher in low-tension (3D) conditions compared to high-tension (2D) conditions. The model predicts that cells interacting with a rBM in 2D that experience high actin tension will form shorter and/or more dynamic membrane protrusions. In contrast, cells that experience low actin tension, as is the case for MECs interacting with a rBM in 3D (3D), are predicted to form longer and more stable membrane protrusions. **(d)** Representative time-lapse confocal microscopy images of MECs stably expressing LifeAct-RFP that were ligated to a rBM in 2D (2D) or 3D (3D). Time in seconds (s) is indicated in each inset; scale bar, 10 μm. **(e)** Bar graph showing quantification of protrusion length in MECs ligated to rBM in 2D, 3D, or in 2D treated with the myosin II inhibitor blebbistatin (2D+ Bleb). (Mean ± s.e.m.; 2D, n= 139; 3D, n=139; 2D+Bleb, n=88). **(f)** Bar graph showing quantification of protrusion residence time in MECs ligated to rBM in 2D (2D), 3D (3D), or in 2D treated with the myosin II inhibitor blebbistatin (2D+ Bleb). (Mean ± s.e.m.; n= 30). **(g)** 30-nm thick computational slices extracted from 3D cryo-ET reconstructions of membrane protrusions in vitrified no-spread MECs cultured on a rBM in 2D in the absence (left) or in the presence of the myosin II inhibitor blebbistatin (right). The position of the cell body is towards the upper right-hand corner (*). Scale bars, 200 nm. Note: under conditions of low cortical actin tension (blebbistatin treatment) the protrusions visualized in these MECs appeared to be highly interdigitated with sharp kinks, consistent with a compliant phenotype. By contrast, the protrusions observed in the untreated samples with higher cortical actin tension were predominantly straight and outwardly projected, suggesting they were stiffer than the protrusions formed in the MECs with lower cortical actin tension (e.g. blebbistatin treated). **(h)** Bar graphs showing quantification of membrane protrusion width in 3D reconstructions of cryo-ET images from no-spread MECs interacting with rBM treated with (low cortical actin tension) and without (high cortical actin tension) blebbistatin (2D + Bleb). (Mean ± s.e.m.; n> 2000). **(i)** Graphical schematic showing how reduced cortical tension influences membrane protrusion phenotype.

Secretory proteins are synthesized in the ER, transported through the Golgi and then to the plasma membrane. Cortical actin physically attaches to the plasma membrane and by so doing influences membrane topology^19^. Membrane topology regulates the binding of curvature-associated proteins that reinforce membrane organization and influence cell signaling and protein secretion or recycling^20^. To address whether the context of ECM engagement facilitates protein secretion by regulating cortical actin tension-dependent plasma membrane topology, we used dynamically triangulated Monte Carlo simulations to explore the morphological conformational space of planar to highly curved membranes^21–23^. The method incorporates (a) thermal fluctuations that play a key role in the self-organization of membranes, (b) the fluid nature of the membrane, and (c) membrane physical environment variables such as tension, osmotic stress, stiffness, and excess area. Using this framework, we quantified the surface deformations induced by membrane proteins as curvature fields, and studied the emergent morphologies of the membrane using isotropic and anisotropic curvature models. This mesoscale model is broadly applicable and appropriate for modeling systems with different tension values. We simulated membrane patches with different excess area (A/A_p_), which is the variable conjugate to the membrane tension and depends on cortical actin tension^23^. Our free energy simulations for the formation of cellular protrusions were computed from the Helmholtz free energy change of the protrusion assembly (ΔF), and were plotted as a function of membrane excess area (Fig. 5b), and thereafter presented as representative cartoons of membrane deformation and protrusion phenotype (Fig. 5c). The model predicted that the protrusions that assemble in the MECs engaging the ECM in 2D that experience high cortical actin tension, herein depicted as being under small excess area (A/A_p_), would be shorter and/or more transient (Fig. 5c). By contrast, the model indicated that the membrane protrusions that assemble in the MECs engaging the ECM in 3D, that experience low cortical actin tension, would be longer and/or more stable. Consistent with these predictions, kymographs of MECs expressing LifeAct-RFP showed that the cells interacting with the laminin-rich rBM in 2D formed highly dynamic, short membrane protrusions, whereas the MECs interacting with the laminin-rich rBM in 3D formed longer and more stable protrusions, quantified as increased protrusion length and residence time (Fig. 5d-f). Causal links between cortical actin tension, membrane protrusion behavior, and membrane topology were demonstrated by showing that reducing myosin activity in the MECs engaging the ECM in 2D (high cortical actin tension) stabilized their plasma membrane protrusions, such that they phenocopied the larger, long-lived plasma membrane protrusions exhibited by MECs engaging the ECM in 3D (low cortical actin tension) (Fig. 5e & f).

We next used correlative light and electron microscopy (CLEM) combined with *in situ* cellular tomography (cryo-ET) to explore causal associations between cortical actin tension, plasma membrane protrusions and plasma membrane topology. CLEM and cryo-ET imaging of the MECs plated on top of rBM-coated EM grids (2D) revealed that the actin-based plasma membrane protrusions were sparsely distributed and predominantly directionally oriented radially away from the cell body (Fig. 5g). By contrast, when these non-spread MECs were treated with blebbistatin to reduce their actomyosin tension, the density of actin-based plasma membrane protrusions was increased and the morphology of the protrusions was modified such that they became heavily interdigitated and convoluted, and sprouted randomly away from the cell body with no preferred directionality (Fig. 5g). Cryo-ET analysis of the membrane protrusions showed a central bundle of actin filaments, with an approximate average distance between the filaments of 117(+/-) nm, consistent with tight bundles mediated by cellular crosslinking molecules such as fimbrin or fascin^24, 25^. This finding indicates that lowering the cortical tension is unlikely to alter the actin crosslinking behavior within the protrusions, Intriguingly, however, statistical analysis of the protrusion widths revealed that the plasma membrane protrusions in the MECs with low cortical tension were on average significantly wider and appear visually more compliant, consistent with their reduced protrusion activity and greater dwell time (Fig. 5g & h). The observations are consistent with the context of ECM engagement could influence ER function and protein secretion by modulating cortical actin tension and plasma membrane protrusions to regulate plasma membrane topology (Fig. 5i).

### Cortical actin tension modulates plasma membrane protein composition

Plasma membrane protrusions are cylindrically shaped structures comprised of a protrusion, an annulus and a basal component that combine to generate negative and positive curvature in the associated membrane (Fig. 6a). While the neck or annulus of the protrusion generates positive membrane curvature, the entire cytoplasmic length of the protrusion compartment generates a negatively curved membrane. Membrane curvature regulates the binding affinity of membrane curvature-associated proteins that can exert profound effects on cell behaviors including endocytosis, protein secretion and recycling, organelle function and actin dynamics^26^. To explore the relationship between cortical actin tension, plasma membrane protrusions and plasma membrane topology, we extended our Monte Carlo model to predict protein recruitment dynamics. We computed the excess chemical potential (or the free energy to add/recruit a protein) to these three spatial regions of the membrane (protrusion, annulus, and basal) as a function of excess membrane area (A/A_p_) (proxy for cortical actin tension) (Fig. 6b). Our calculations revealed that a lower value of the excess chemical potential signifies the favorable recruitment of a protein to a given membrane location and curvature to enhance its local protein density. As shown in Fig. 6b, our computational analysis revealed that the excess chemical potential of negative curvature-binding protein domains is preferentially localized to the cytosolic site of the membrane protrusions, and predicted that this protein domain binding behavior will segregate more favorably under conditions of low cortical actin tension; 3D. Our energetics calculations further predicted that conditions in which cell tension is low (MECs engaging the ECM in 3D), the recruitment of positive curvature-inducing proteins would be hindered (Fig. S6a).

**Figure 6.**
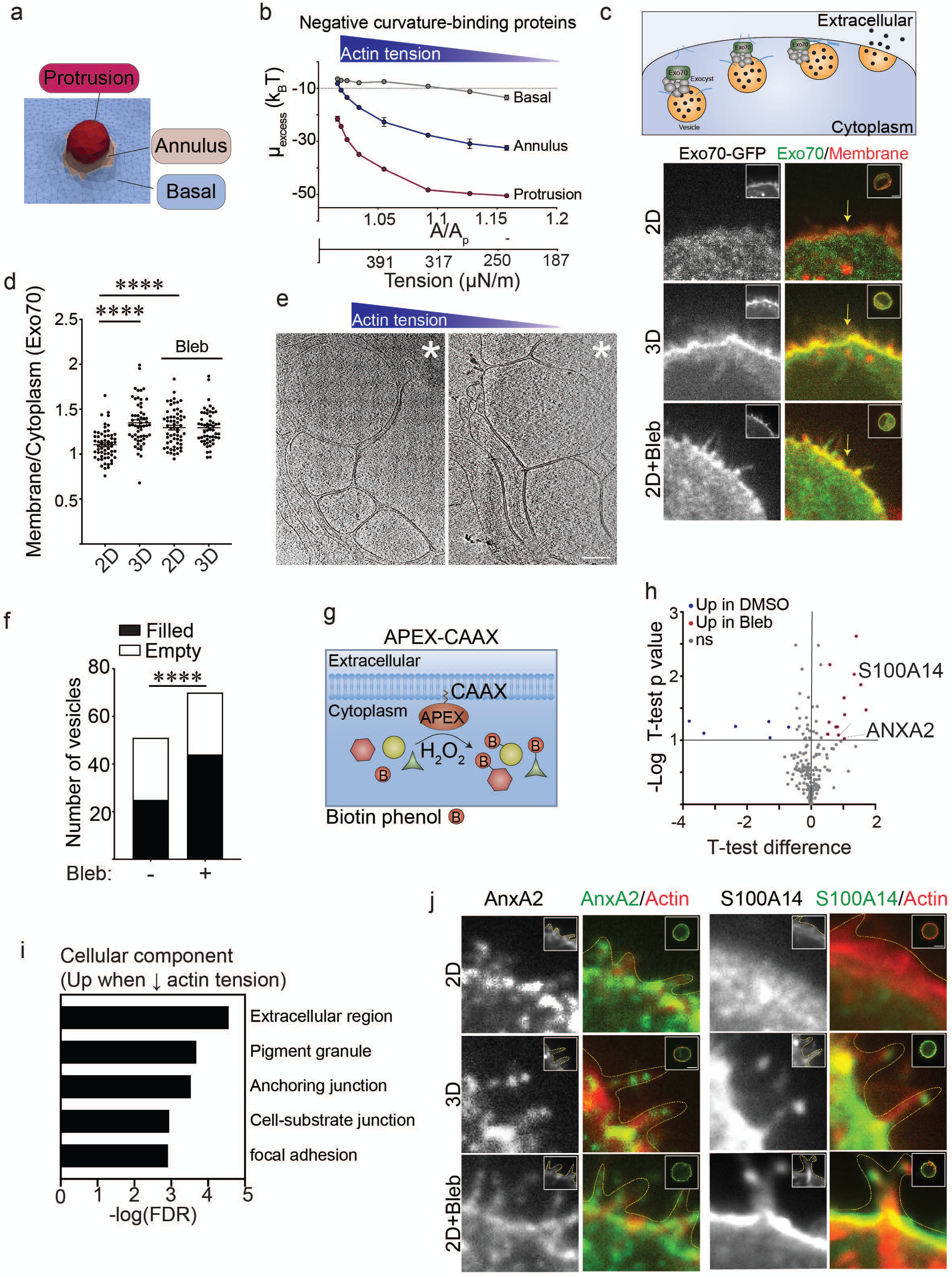
Cortical actin tension modulates plasma membrane protein association. **(a)** Cartoon showing spatial regions associated with plasma membrane actin-associated projections including the basal domain, the annulus and the protrusion. **(b)** Line graph illustrating excess chemical potential required to recruit negative curvature sensing domain proteins to the basal, annulus, and protrusion region of the plasma membrane as a function of excess membrane area (A/A_p_). **(c)** (Top) Cartoon showing how exocyst components including Exo70 tether vesicles during exocytosis. Representative fluorescence microscopy images of MECs stably expressing EGFP-Exo70 and farnesylated mCherry (membrane tag; insets in the left column) that were ligated to rBM in 2D (2D), 3D (3D), and in 2D treated with blebbistatin (2D + Bleb) to reduce cortical actin tension. Fluorescent images of whole cells are shown (inset; right) and arrows point to the cell edge. Scale bar, 10 μm. **(d)** Bar graphs showing quantification of EGFP-Exo70 localization to the plasma membrane in MECs ligated to rBM in 2D and 3D treated with (Bleb) and without blebbistatin to reduce myosin II activity. Membrane localization of EGFP-Exo70 was quantified as a ratio of plasma membrane to cytoplasm fluorescence intensity (Mean ± s.e.m.; n=50 cells per condition). **(e)** 30-nm thick computational slices extracted from 3D cryo-ET reconstructions of vitrified MECs cultured on rBM in 2D treated with (right panel) or without (left panel) blebbistatin to reduce myosin II activity and cortical actin tension, showing subplasmalemmal regions enriched with membrane-engulfed compartments. The position of the cell body is towards the upper right-hand corners (*). Scale bars, 200 nm. **(f)** Quantitative analysis of the membrane-engulfed compartment classified as either ‘filled’ (black) or ‘empty’ (white), based on the presence and absence of recognizable macromolecules. Bar graphs show quantification of the number of vesicles proximal to the plasma membrane. **(g)** Schematic diagram of proximity-based biotinylation assay used to label proteins localized to the plasma membrane under different ECM ligation conditions. Expression of a recombinant APEX2-CAAX probe within cells facilitated biotin labelling of proteins proximal to the plasma membrane upon addition of biotin-phenol and H_2_O_2_. **(h)** Volcano plot showing differentially associated proteins in MECs ligated to rBM in 2D (blue; log 2> 0.5; high cortical actin tension) compared to those ligating a rBM in 2D treated with blebbistatin to reduce myosin II activity (red; log 2> 0.5; low cortical actin tension). Differentially expressed genes with an adjusted p-value<0.1 and log 2 (fold change) > 0.5 are shown (n=2 biologically independent experiments). **(i)** GO Cellular Component analysis of proteins enriched at the plasma membrane when cortical tension was reduced. **(j)** Representative immunofluorescence microscopy images of MECs ligated to rBM in 2D (2D), 3D (3D), or in 2D treated with blebbistatin (2D+ Bleb) to reduce myosin II activity, and immunostained with antibodies targeting Annexin A2 (AnxA2) or S100A14. Cellular F-actin was counterstained using Alexa569-labelled phalloidin (insets in the first and third columns) and the cell edge is marked with a yellow dashed line. Fluorescent images shown are of whole cells (insets in the AnxA2/Actin and S100A14/Actin columns). Scale bar, 10 μm.

Given that Exo70 is a negative curvature binding/inducing protein that is also a component of the exocyst complex targeting secretory vesicles to the plasma membrane (Fig. 6c; top panel), we predicted that Exo70 could be a candidate protein regulated by cortical actin tension to facilitate secretory protein trafficking^27^. Consistent with our energetics model prediction, EGFP-tagged Exo70 showed greater plasma membrane localization in the MECs engaging the ECM in 3D (low actin tension) as revealed by a strong co-localization of the EGFP-Exo70 with membrane-tagged farnesylated mCherry as compared to minimal membrane associations observed in the MECs engaging the ECM in 2D (Fig. 6c & d). Reducing myosin activity in the MECs engaging the ECM in 2D (blebbistatin) to recapitulate the membrane protrusion behavior of the MECs engaging the ECM in 3D, significantly enhanced the level of plasma membrane-associated Exo70, implicating cortical actin tension in this plasma membrane phenotype (compare Fig. 6c & d; compare 2D top to 2D+Bleb bottom panels). Indeed, statistical analysis of the 3D cryo-ET imaging reconstructions containing membrane-enclosed compartments revealed that the cells with reduced cortical tension (3D) had more membrane-enclosed compartments proximal to the plasma membrane region than the cells with high cortical tension (2D) (Fig. 6e & f). Moreover, the analysis revealed that a higher fraction of the membrane compartments in the MECs with reduced actin tension were enriched with macromolecular content, whereas a higher fraction of vesicle-like structures adjacent to the plasma membrane in the MECs with high actin tension appeared emptier (Fig. 6e & f), suggesting defective secretory pathway cargo loading. Consistently, RT-qPCR analysis demonstrated that the MECs with high cortical actin tension expressed lower levels of SEC61B, an important protein required for protein insertion into the ER (Fig. 5a). The data suggest cortical actin tension could regulate secretory protein trafficking to the plasma membrane by regulating plasma membrane protrusion activity and topology. This possibility accords with the RNA-seq data comparing gene expression in the MECs engaging the ECM in 2D versus 3D that revealed a significant increase in the expression of several ER protein insertion regulators and secretory pathway regulators in the MECs with low cortical actin tension (ligating ECM in 3D)(ligating ECM in 2D) (Fig. 1k).

To explore the possibility that cortical actin tension modulates plasma membrane protein composition, we used an unbiased proximity-based biotinylation assay to identify a broader repertoire of cell cortex-associated proteins whose localization is regulated by myosin activity. Using membrane-targeting ascorbate peroxidase (APEX2-CAAX) as a bait, we mapped the proteomic landscape at the actin cortex in MECs with high and low myosin activity (Fig. 6g; MECs in 2D with and without blebbistatin treatment). We used APEX2-CAAX to mark the cellular cortex (Fig. S6a) and immunoblotting to verify APEX biotinylation efficiency (Fig. S6b). Quantitative mass spectrometry analysis identified 13 candidate proteins whose plasma membrane associations were modulated by myosin activity (Fig. 6h & i). Gene enrichment analysis revealed that these tension-sensitive membrane-associated proteins were predominantly localized at either the extracellular region of the cell, at cellular vesicles, or at the cell-substrate interface (Fig. 6i). Immunofluorescent staining verified that the localization of the APEX2-CAAX identified protein AnxA2 was enriched at the plasma membrane as well as within plasma membrane protrusions in the MECs with low cortical actin tension (Fig. 6j; compare AnxA2 2D to 3D and 2D+Bleb panels). AnxA2 can form heterodimeric structures with S100 family members, which are extracellular secreted proteins, known to form a bridge between two membranes to facilitate vesicle docking during exocytosis^28^. Consistently, immunofluorescence staining for S100A14 showed abundant plasma membrane and membrane protrusion localization in the MECs with low cortical actin tension (Fig. 6j; compare S100A14 2D to 3D and 2D+Bleb panels).

The increased AnxA2 and Exo70 plasma membrane localization we observed is not only consistent with the Molecular Dynamic Simulations that predicted the plasma membrane binding of negative curvature-binding proteins, such as AnxA2 and Exo70, but also suggested that this binding could induce and stabilize negative plasma membrane curvature^27, 29^. Accordingly, the findings suggest that cortical actin tension regulates membrane protrusion activity and plasma membrane topology to modulate the spatial distribution of negative curvature-binding proteins implicated in secretory protein trafficking.

### Cortical actin tension regulates MEC spheroid phenotype

Our results thus far implied that how a cell ligates its ECM has a profound effect on ER function and possibly protein secretion. We implicated filamin-dependent cortical actin tension as a key regulator of this ECM ligation mediated ER homeostasis phenotype. Unresolved or prolonged ER stress responses compromise cell viability, and impeding secretory protein trafficking can severely compromises ER function^30^. Therefore, we examined whether the context of ECM ligation regulates MEC viability at the single cell level and influences spheroid stress resilience and if this is mediated through tuning cortical actin tension. To begin with, calcein/ethidium homodimer live/dead staining revealed the single non-spread MECs with high cortical actin tension that engaged the ECM in 2D had significantly reduced cell viability as compared to non-spread MECs engaging the ECM in 3D with low cortical actin tension (Fig. 7a; left). However, and importantly, the viability of the non-spread MECs engaging the ECM in 2D could be significantly extended if their myosin activity was reduced by blebbistatin treatment (Fig. 7a; right). Consistently, reducing filamin using shRNA simultaneously enhanced cellular viability in the non-spread MECs engaging the laminin-rich rBM in 2D (Fig. 7b). Manipulating cortical actin tension either by inhibiting myosin activity or through filamin knockdown had no impact on cell adhesion or cell spreading implying cell viability regulation occurred through other mechanisms. Indeed, decreasing cellular filamin significantly ameliorated ER stress signaling in the MECs interacting with the ECM in 2D as indicated by ATF3 expression level (Fig. 3f). Consistently, knocking down the negative membrane binding protein Exo70, which has been implicated in protein secretion and membrane topology regulation, compromised the viability of the MECs interacting with the laminin-rich rBM in 3D, again without altering cell spreading or cell-ECM adhesions (Fig. 7c). The data therefore suggest that filamin-dependent cortical actin tension modulates ER function to regulate MEC viability.

**Figure 7.**
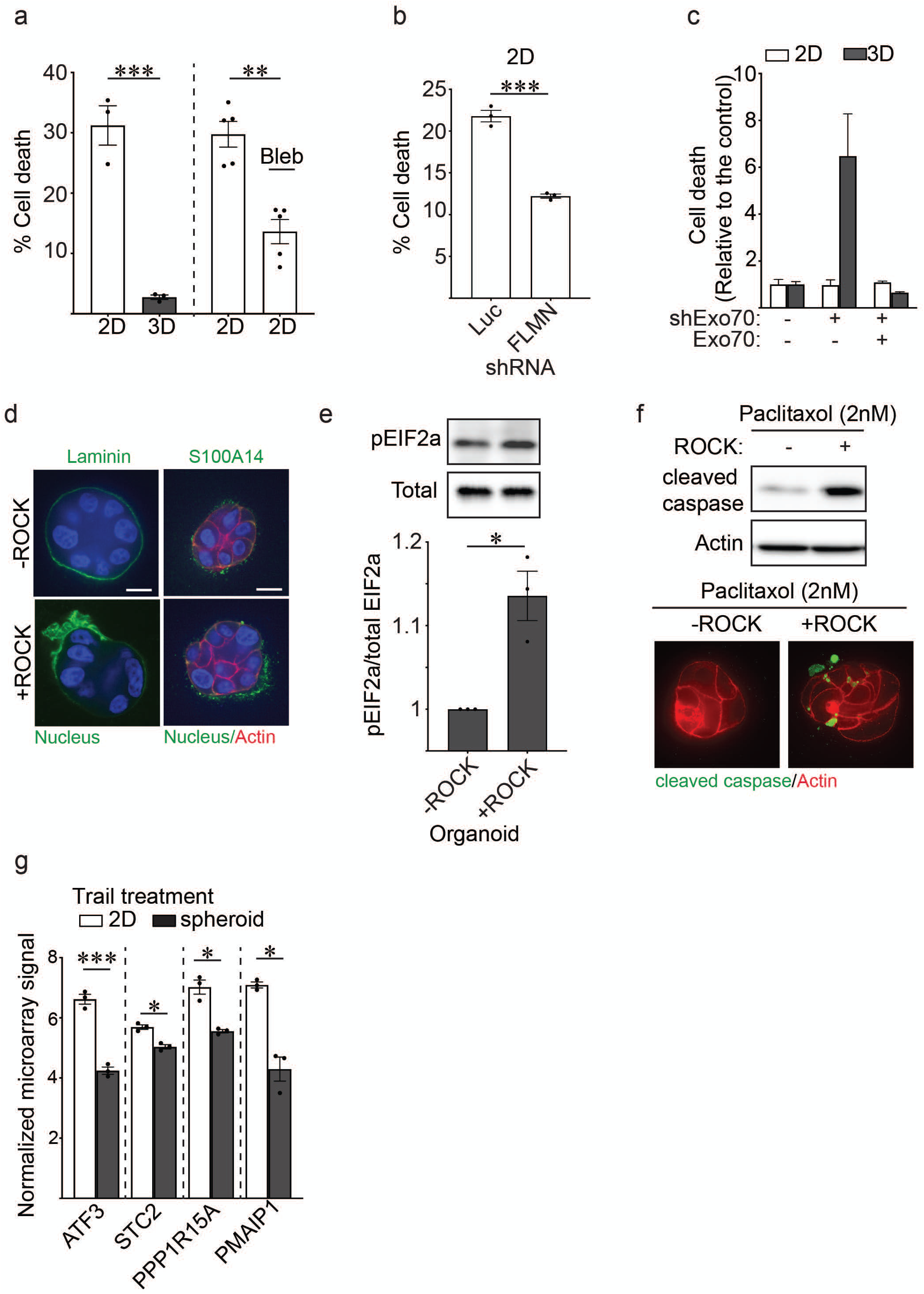
Cortical actin tension regulates MEC spheroid phenotype. **(a)** Graphs showing percent cell death measured in MECs ligated to rBM in 2D (high cortical actin tension) and 3D (low cortical actin tension) and in 2D treated with blebbistatin to reduce myosin II activity (low cortical actin tension;2D+Bleb) 48 hr post-plating as quantified using calcein AM and ethidium homodimer staining. (Mean ± s.e.m.; 2D vs 3D, n = 3; Blebbistatin treatment, n = 5). **(b)** Bar graphs showing percent cell death quantified using calcein AM and ethidium homodimer staining 48hr post-plating in MECs stably expressing shRNA that targets filamin (FLMN) or luciferase (Luc) that were ligated to rBM in 2D. (Mean ± s.e.m.; n=3). **(c)** Bar graphs showing percent cell death quantified using calcein AM and ethidium homodimer staining 48hr post-plating in MECs stably expressing shRNA that targets Exo70 or luciferase (Luc). (Mean ± s.e.m.; n=3). **(d)** Spheroids with or without inducible, constitutively active ROCK expression were stained with antibodies for laminin or S100A14 (green). The actin and nucleus were counterstained with phalloidin (red) and DAPI (blue), respectively. Scale bar, 10 μm. **(e)** Bar graphs showing the level of phosphorylated EIF2a (pEIF2a) and total EIF2a in MEC spheroids with and without ROCK activation, evaluated via immunoblotting and quantified as a fraction of total EIF2a protein (Mean ± s.e.m.; n=3). **(f)** Spheroids with or without inducible, constitutively active ROCK expression were treated with Paclitaxol (2nM) and stained with antibodies for cleaved caspase-3 (green). Actin filaments were counterstained with phalloidin (red). The level of cleaved caspase-3 was also evaluated via immunoblotting. (g) Bar graphs show the relative mRNA level as quantified by microarray analysis of ER stress response genes (ATF3, STC2, PPP1R15A, and PMAIP1) in TRAIL treated MECs plated as monolayers on a rigid rBM (2D; high cortical actin tension) or as spheroids (3D; low cortical actin tension). (Mean ± s.e.m.; n = 3).

We next explored if the reduction in cortical actin tension induced by ligation of an ECM in 3D could account for polarized protein secretion and the stress resilience phenotype of MEC spheroids embedded within a laminin-rich rBM. We ectopically increased actomyosin tension in preassembled MEC spheroids embedded within a laminin-rich rBM by expressing an inducible, constitutively active ROCK and then monitored effects on endogenous basement membrane deposition, polarized protein secretion and resistance to exogenous stress. Consistent with a causal association between cortical actin tension and ER homeostasis, confocal imaging revealed that increasing cortical tension through activated ROCK compromised the assembly of an endogenous, polarized basement membrane as indicated by disorganized laminin-111 IF staining (Fig. 7d). Furthermore, elevating ROCK activity also disrupted the level and membrane association of S100A14 secreted protein (Fig. 7d). Increasing cortical tension in the spheroids by activating ROCK also increased levels of phosphorylated EIF2a (pEIF2a) which together with the compromised protein secretion imply high cortical actin tension increases ER stress (Fig. 7e). To further explore the relationship between actin tension and ER function we monitored the chemotherapy response of MEC spheroids embedded within the laminin-rich rBM. Results showed that increasing actomyosin tension by expressing the activated ROCK sensitized the MEC spheroids to the chemotherapy agent Paclitaxol, as quantified by increased levels of cleaved caspase-3 (Fig. 7f). Similarly, bioinformatics analysis revealed that many genes significantly induced in the TRAIL-treated MEC monolayers interacting with the laminin-rich rBM in 2D, with high cortical actin tension, as compared to those MEC spheroids interacting with the laminin-rich rBM in 3D with low cortical actin tension were genes implicated in PERK-dependent ER stress regulation including ATF3, STC2, PPP1R15A (Fig. 7g). The findings suggest high filamin-dependent cortical actin tension compromises ER homeostasis to perturb protein secretion and activate an ER stress response activity and implicate PERK-dependent signaling in this phenotype.

## Discussion

Our studies underscore the importance of cellular context in regulating organelle homeostasis and tissue phenotype. Our data demonstrate that the context of cell-ECM ligation regulates ER function including ER stress response signaling and secretory protein trafficking and does so by modulating cortical actin tension. Our findings revealed that cortical actin and actin-binding molecules, which regulate cellular tension, are critical effectors of the tissue-like behavior of cells interacting with an ECM in 3D. Our observations that ECM topology affects ER homeostasis are consistent with prior findings showing stress-resilient, enhanced protein secretion and deposition by spheroids and organoids embedded within a rBM (3D) and offer a plausible explanation for why these tissue-like structures exhibit such profound treatment resistance and are such useful models for drug testing. These findings also clarify why organoids cultured within a rBM (3D) are able to faithfully reconstitute self-organization phenotypes and tissue behaviors such as branching morphogenesis and tubulogenesis.

Our results revealed that secretory protein trafficking is perturbed when cortical actin tension is elevated. Cortical actin tension could regulate secretory protein trafficking on multiple levels. Indeed, secretory protein synthesis and trafficking is impaired during ER stress^18^, suggesting that cortical actin tension influences protein secretion, in part, by modulating ER homeostasis. Surprisingly, we found that actin tension can also affect plasma membrane topology to alter the localization/composition of targeting secretory effectors at the plasma membrane. In particular, membrane protrusion formation and the recruitment of negative curvature binding proteins including Exo70 and ANXA2, which serve important functions in membrane trafficking and protein secretion, require low cortical actin tension and are inhibited by high actin tension^28, 31^.

Our studies showed how ER function is sensitive to cortical actin tension and suggest that the ER harbors mechanosensitive elements. Indeed, recent evidence suggests that numerous calcium channels localized to the ER (e.g. Pannexin-1 and Piezo) can be activated by extracellular mechanical stimuli such as uniaxial stretching, focused ultrasound and laser-tweezer-traction at the plasma membrane^32–34^. We consistently found that cells ligated to an ECM in 2D generate high actin tension and simultaneously develop dysregulated calcium homeostasis. These finding raise the intriguing possibility that the high actin tension in cells interacting with an ECM in 2D aberrantly activates mechanosensitive ER calcium channels, ultimately compromising calcium homeostasis, ER function, and cell viability.

So how does the actin cortex influence ER function? Our data suggest that the actin crosslinker filamin is an important regulator of this process. Filamin is not only an actin crosslinker required for the maintenance of cortical actin tension but also a critical adaptor protein that operates at ER-PM contact sites to facilitate SOCE^13^. Given that we did not see an increase of intracellular calcium levels or SOCE under high actin tension (2D; high filamin expression), we speculate that ER-PM contact formation in cells ligated to an ECM in 2D serves as a feedback mechanism important for the re-establishment of calcium homeostasis. Filamin-mediated actin crosslinking likely influences ER function via one of the following two mechanisms. First, filamin is required to maintain actin cortex integrity and tension. High filamin-mediated cortical tension may activate mechanosensitive calcium channels in the ER to modulate calcium homeostasis and therefore ER function. Alternatively, given that high actin tension impaired exo70 recruitment to the plasma membrane, a stiffer filamin-crosslinked actin cortex in 2D conditions may be less amenable to secretory activity, possibly leading to aberrant accumulation of proteins in the ER and activation of ER stress responses. Regardless of how filamin regulates the ER, we present compelling evidence that cortical actin composition and tension are crucial for ER homeostasis. Our results also broach the idea that cellular conditions which increase cortical actin tension, including a pathologically stiffened ECM or elevated actomyosin tension, would similarly activate ER stress response signaling in cells. Indeed, tumors that develop within a stiffened fibrotic ECM and express oncogenic Ras, which increases ROCK activity and actomyosin tension, demonstrate compromised ER stress regulation^32, 35^.

External stresses such as hypoxia and nutrient deprivation trigger ER stress. For example, suspended cells that are devoid of ECM ligation also activate PERK-dependent phosphorylation of EIF2a and selectively upregulate ATF4 expression^36^. Notably, our studies demonstrate that actin tension is necessary and sufficient to modulate ER function since (i) ER stress signaling can be alleviated by agents that reduce cortical actin tension without affecting cell-ECM ligation (2D + Blebbistatin), whereas (ii) ROCK-dependent induction of cortical actin tension promoted ER stress signaling, even in the presence of 3D ligation. These findings raise the possibility that suspended cells devoid of ECM ligation also have altered actin cortex composition or mechanics, contributing to the ER stress phenotypes observed under these conditions^36^.

## Acknowledgments

This work was supported by the following grants: US National Institutes of Health NCI grants R35 CA242447-01A1; R01CA222508-01 ; U01 CA202241 and U01 CA250044-01A1; and BCRF A132292 (V.M.W.); Canadian Institutes of Health Research Postdoctoral Fellowship (F.B.K.); National Science Foundation Graduate Research Fellowship (G.O. and A.F.L); U01 grant CA202241 (C.S.); NIH R01GM1341 (S.D); NCI1U01CA202123 (Y.L.H and M.G.); NIH/NIGMS DP2OD022552 (H.H.H. and A.P.W.); American Association for Cancer Research Basic Cancer Research Fellowship (J.J.N.); Banting Postdoctoral Fellowship from the Government of Canada (A.M.L.); NIH Program Project Grant P01 GM121203 (N.V.), R01 AI132378 (D.H. & N.V.), NIH Grants S10-OD012372 (D.H.), S10-OD026926 (D.H.), P01-GM098412-S1 (D.H.) and Pew Charitable Trust 864K625 (D.H.) funded the purchase of the Titan Krios TEM, Falcon 3EC DDD, and Eagle CMOS (Thermo Fisher Scientific) and cryo-CLEM equipment (Leica Microsystems). NIH R35 GM141832 (W.G.); NIH CA227550 and CA193417 (R.R. and R.W.T.)

## Author Contributions

V.M.W. and F.B.K. conceived the study and designed the experiments. R.R. and R.W.T. performed computational modeling analysis. C.S. performed bioinformatics analysis of the RNA-seq and microarray data. D.H. and G.G. designed the CLEM, cryo-TEM and cryo-ET workflow employed here, with assistance from M.F.S., F.B.K and V.M.W. N.V. pursued the reconstructions, image processing, data mining, and statistical analysis of the CLEM and cryo-ET data with feedback from G.G. M.F.S and D.H. G.O., A.F.L. and S.D. optimized and performed the laser ablation experiments. G.O., J.J.N. and A.M.L. performed biochemical and cell biological experiments. Y.L.H. and M. G. performed microrheology measurement. H. H. H. and A.P.W. optimized the mass spectrometry setup, performed mass spectrometry and analyzed the resulting data. V.V. provided microfabrication advices and reagents. V.V., D.M.B, and W.G. provided scientific suggestions on the projects. V.M.W. and F.B.K. wrote the manuscript with input from all authors.

## Data Availability

The authors declare that all data supporting the findings of this study are available within the paper and its supplementary information files.

## Materials and Methods

### Cell culture

MCF10A cells were cultured in DMEM/F12 Ham’s mixture (Invitrogen) supplemented with 5% horse serum (Invitrogen), 20 ng/ml EGF (PeproTech), 10 µg/ml insulin (Sigma), 100 ng/ml cholera toxin (Sigma), and 0.5 mg/ml hydrocortisone (Sigma). HEK293T cells (ATCC, CRL-3216) were cultured in DMEM (Genesee Scientific) supplemented with 10% FBS (Gemini Bio-Products). HMT-3522 S-1 MECs were cultured in chemically defined medium as previously described^50^. All cells and derivative stable cell lines were maintained at 37 °C and 5% CO_2_ and were routinely tested for mycoplasma contamination.

### Antibodies and chemicals

Reagents for immunofluorescence staining: mouse anti-filamin (Millipore), rabbit anti-Annexin A2 (Cell Signaling), rabbit anti-S100A14 (Sigma), CF-dye phalloidin conjugates (Biotium). Antibodies for immunoblotting: rabbit anti-V5 (Cell Signaling), rabbit anti-GAPDH (Cell Signaling), rabbit anti-phospho-EIF2a (Cell Signaling), rabbit anti-EIF2a (Cell Signaling), rabbit cleaved caspase-3 antibody (Cell Signaling), rabbit Esyt1 antibody (ABclonal), HRP-conjugated goat anti-rabbit (Advansta), and HRP-conjugated goat anti-mouse (Advansta). Other reagents include: recombinant basement membrane (rBM; BD Biosciences), Cultrex 3D Culture Matrix Laminin I (R&D systems), blebbistatin (Sigma), polyethylenimine (Polyplus Transfection), doxycycline (Sigma), Isopropyl β-d-1-thiogalactopyranoside (IPTG) (Sigma), 4-hydroxytamoxifen (Sigma), Biotin-phenol (Sigma).

### 2D and 3D culture setup

2D monolayer cultures were plated on rBM-laminated glass coverslips overnight. For fully embedded 3D culture, cells were resuspended in an ice-chilled solution (composed of 50% Matrigel and 50% growth media) and immediately placed at 37 °C for matrix gelation for 1hr. Regular growth media were supplemented to the well 45 min after Matrigel polymerization. Polyacrylamide (PA) gels were prepared as described previously^51^. Briefly, compliant PA gels were polymerized on silanized coverslips and were either functionalized with Matrigel (50 µg/ml) or laminin (40 µg/ml) overnight. PA gels were washed three times and equilibrated overnight at 37 °C prior to cell seeding. Cells that remained unattached 1hr post-seeding were removed by replacing the supernatant with regular growth media. In 2D conditions, the supernatant was replaced with fresh media, whereas in 3D conditions, the supernatant was replaced with fresh media supplemented with 2% Matrigel or 100 µg/ml laminin, which generates a 3D scaffold around cells. Micropatterned surfaces were prepared as described previously^52^. Briefly, a thin film of round perforations (microwells) 10 µm in diameter were fabricated and adhered to tissue culture-treated coverslips. Alexa-647 conjugated laminin-111 (40 µg/ml) was used to coat the surface overnight at 4 °C. After two washes with PBS, the thin microwell film was subsequently removed. PEG-PLL (100 µg/ml) was used to passivate the unpatterned glass surface for 1hr. The fluorescent signal of laminin-111 on the coverslips was used to ensure proper patterning prior to cell plating. Cells were trypsinized and incubated on the micropatterned surface for 15 min. For 2D conditions, unattached cells were removed by aspiration and replaced with fresh media. In 3D conditions, unattached cells were similarly removed and replaced with fresh media supplemented with 100 µg/ml of laminin.

### Microarray experiments

Total RNA for microarray analysis was extracted from three independent culture preparations using TRIzol^TM^ (Life Technologies) and RNA was purified using a RNeasy Mini Kit and a DNase treatment (Qiagen). To prevent apoptosis induction, organoid cultures were pretreated with caspase inhibitors DEVD-CHO and Ac-IETD-CHO (1 µM, respectively) at least 2hr before stimulation and inhibition was maintained throughout the duration of TRAIL treatment. Gene expression analysis was performed on an Affymetrix GeneChipTM Human Genome U133A 2.0 platform containing 22,283 probes according to the manufacturer’s protocol. Biotinylated cRNA was produced from total RNA from each sample. After hybridization, washing and staining, arrays were scanned using a confocal scanner. The hybridization intensity data was processed using the GeneChip Operating software. Affymetrix.cel files (probe intensity files) were processed with the Transcriptome Analysis Console (TAC) software. A filtering criterion (P < 0.05 by empirical Bayes test, fold-change > 2.0-fold) was used to select differentially expressed genes within a comparison group. Gene ontology analysis was performed using GAGE on RMA-normalized microarray data with the ’affy_hg_u133a’ and ’name_1006’ attributes from the ’ensembl’ biomaRt. The microarray data can be download from NCBI Gene Expression Omnibus website (GEO: GSE138900).

### RNA purification, RNA-seq library preparation, sequencing, and analysis

RNA was isolated using TRIzol^TM^ (Life Technologies) followed by chloroform extraction. RNA pellets were washed three times in 75% isopropanol and resuspended in RNAse-free water. The integrity of total RNA was examined by formaldehyde-agarose gel electrophoresis. RNA-seq libraries were constructed using a combination of KAPA mRNA HyperPrep Kit (Illumina) and NEBNext RNA Library Prep Kit (New England Biolabs) protocols. The poly-A-containing mRNA (500 ng) were purified using oligo(dT) beads and fragmented into 200-300bp fragments using heat and magnesium. The cleaved RNA fragments were reversed transcribed into first strand cDNA using random primers. Second strand synthesis and an A-tailing step were performed according to Kapa kit instructions. NEBNextAdaptor was ligated to dA-tailed DNA and the end products were gel purified to remove excess unligated adaptors. USER enzyme was used to cleave at the uracil incorporation sites followed by library amplification using NEBNext index primers for multiplexing. At the final stage, the cDNA library was again purified using gel electrophoresis. Libraries were quantified on a Bioanalyzer and sequenced via HiSeq4000 (Illumina) at the Vincent J. Coates Genomics Sequencing Laboratory, University of California, Berkeley. We performed 50-bp single-end sequencing on all samples and sequencing reads were aligned to the Gencode human genome primary assembly (v31) using STAR (v2.7.1a). Aligned reads were counted for each gene using the analyzeRepeats.pl function in HOMER (v4.9.1). Low expression genes were filtered out if they did not have at least one count per million (CPM) in at least three samples and normalized using the calcNormFactors function in edgeR with the “TMM” method. Clustering was performed on log2(CPM) values using Limma’s plotMDS function. Differential expression analysis was done using Limma-Voom. Gene ontology analysis was applied to log2(CPM) values using GAGE with the ’ensembl_gene_id_version’ and ’name_1006’ attributes from the ’ensembl’ mart on biomaRt.

### Quantitative RT-PCR

Total RNA was harvested using a TRIzol^TM^ (Life Technologies) method as described above. Samples were reverse transcribed using random hexamer primers. PerfeCTa SYBR Green FastMix (Quantibio) was added to each cDNA and the sequences of primers used in this study are listed below: GAPDH-For: CAGCCTCAAGATCATCAGCA; GAPDH-Rev: TGTGGTCATGAGTCCTTCCA; SEC61B-For: TCATCTCCAATATGCCTGGTC; SEC61B-Rev: TTTGAGCCCAGGTGAATCTT; SERP1-For: GGACCCTGGTTATTGGCTCT; SERP1-Rev: CATTCTGAGGCAGGATGCTT. ATF3-For: CGCTGGAATCAGTCACTGTCAG; ATF3-Rev: CTTGTTTCGGCACTTTGCAGCTG; STC2-For: GGGAATGCTACCTCAAGCAC; STC2-Rev: CACAGGTCAGCAGCAAGTTC. Thermal cycling conditions were 10 min at 95 °C, followed by 40 cycles of 15s at 95°C and 45s at 65°C. Melting curve analysis was performed at the end of each PCR to verify the specificity of each primer. Relative mRNA expression was determined by the ΔΔCT method, normalized to GAPDH.

### Generation of expression constructs

A detailed description of cDNA constructs and their construction is provided in Supplementary Note 1.

### Viral packaging, infection and selection and transfection

To generate retrovirus (pBabe and pBMN vectors), Phoenix cells were seeded and transfected with the indicated vectors using polyethylenimine (PEI). For lentivirus packaging (pLV or IPTG-inducible pLKO-shRNA vectors), HEK293T cells were seeded and co-transfected with psPAX2 and pMD2.G and indicated lentiviral vectors using PEI. Virus-containing supernatant was collected 48 hr post transfection and the supernatant was centrifuged twice to remove residual Phoenix or HEK293T cells in suspension. The supernatant was supplemented with 8 µg/ml of polybrene prior to infection. Cells were infected with virus overnight, recovered from viral infection for 24hr followed by either puromycin or G418 selection until all uninfected control cells were eliminated by the antibiotic. To generate doxycycline-inducible filamin-overexpressing MECs, cells stably expressing the synthetic reverse Tet transcriptional transactivator rtTAs-M2 were transiently co-transfected with a transposon pPB-Puro-Tet construct expressing filamin and a construct expressing a hyperactive piggyBac transposase (pCMV-HAhyPBase) using Lipofectamine 3000 Reagents. Cells were allowed to recover from transfection for 24hr prior to puromycin selection. Doxycycline (0.5 µg/ml) was used to induce filamin overexpression in MECs for 24hr prior to experiments. To generate filamin knockdown cells, MECs were infected with virus particles packaged with constructs expressing either Luciferase or filamin shRNAs. The expression of luciferase or filamin shRNAs in MECs was induced using IPTG (100 µM) for 5 days prior to experiments.

### Immunofluorescence

Cells on compliant PA gels or micropatterned surfaces were rinsed twice with PBS prior to fixation using 4% formaldehyde for 15 minutes. Cells were subsequently permeabilized with 0.2% Triton X-100, quenched in 10 mM glycine/PBS, blocked in 10% goat serum/1% BSA, and incubated with primary and secondary antibodies in 1% goat serum. Secondary antibodies were conjugated to either Alexa 488, Alexa 555 or Alexa 647. Immmunofluorescence and live cell imaging was performed on an inverted microscope (Eclipse Ti-E; Nikon) with spinning disk confocal (CSU-X1; Yokogawa Electric Corporation), 405 nm, 488 nm, 561, 635 nm lasers; an Apo TIRF 100XNA 1.49 objective; electronic shutters; a charge-coupled device camera (Clara; Andor) and controlled by Metamorph. Fully embedded spheroid staining was conducted as above except that spheroids were fixed using 4% formaldehyde supplemented with 0.5% glutaraldehyde to prevent rBM depolymerization during fixation.

### Intracellular calcium imaging

MECs were loaded with 2 µM Fura-AM in extracellular buffer (125 mM NaCl, 5 mM KCl, 1.5 mM MgCl_2_, 20 mM HEPES, 10 mM glucose, 1mM CaCl_2_) for 30 min at room temperature. Loaded cells were washed twice with extracellular buffer twice and incubated for another 30 min before imaging. Fura-AM fluorescence is acquired by illuminating cells with analternating 340/380nm light every 5s and fluorescent intensity is collected at 510nm. Cells were imaged briefly and treated with 3 mM EGTA and 2 µM Thapsigargin in calcium-free HBSS to stimulate ER Ca^2+^ release and chelating extracellular Ca^2+^. After 15 min of imaging, 10 mM CaCl_2_ was added back to the cells to allow for calcium influx.

### Traction force microscopy

Cells were plated sparsely on rBM-laminated compliant 400Pa PA gels containing 100 nm fluorescent beads. The gel was mounted on a microscope chamber to maintain 37 °C and 5% CO_2_. Phase contrast images were taken to record cell position and fluorescent images of beads embedded in the gel just below the cells were taken to determine gel deformation. Images of MECs were collected before and after 0.5% SDS treatment using a Nikon Inverted Eclipse TE300 microscope and a Photometric Cool Snap HQ camera (Roper Scientific). Images were exported to ImageJ and aligned using the StackReg plugin (NIH). The bead displacement field and the force field were reconstructed using Iterative Particle Image Velocimetry (PIV) and Fourier Transform Traction Cytometry (FTTC) plugins from Image J, respectively. The freely available package of traction force microscopy software is available at https://sites.google.com/site/qingzongtseng/tfm.

### Atomic force microscopy

All AFM measurements were performed using a MFP3D-BIO inverted optical AFM (Asylum Research, Santa Barbara, CA) mounted on a Nikon TE200-U inverted microscope (Melville, NY) and placed on a vibration-isolation table (Herzan TS-150). A 2-µm beaded tip attached to a silicon nitride cantilever (Asylum Research, Santa Barbara, CA) was used for indentation. The spring constant of the cantilever was 0.06 N/m. For each session, cantilevers were calibrated using the thermal fluctuation method^53^. AFM force maps were performed on 40 x 40 µm fields and obtained as a 12 x 12 raster series of indentation. Elastic moduli measurement was derived from the force curves obtained utilizing the FMAP function of the Igor Pro v. 6.22A (WaveMetrics, Lake Oswego, OR) supplied by Asylum Research. Cells were assumed to be incompressible and a Poisson’s ratio of 0.5 was used in the calculation of the Young’s elastic modulus. 2D measurements were conducted on MECs plated on a laminin-conjugated micropatterned surface with media overlay. 3D measurements were obtained using MECs incubated overnight with media containing laminin-111 (100 µg/ml).

### Optical tweezer measurement

Mechanical properties of the cytoplasm were measured as described previously^29^.Briefly, 0.5 μm latex beads were endocytosed by cells before they were seeded on compliant PA gels. Upon attachment, cells were cultured in either 2D or 3D conditions, as described above. The laser beam (10W, 1064 nm) was focused through a series of Keplerian beam expanders and a 100X oil immersion objective. A high-resolution quadrant detector was used for position detection. The endocytosed bead was dragged at a constant velocity of 0.5 μm/s by the optical trap, and the force displacement curve of the local cytoplasm was recorded. The cytoplasmic modulus was calculated based on the slope in the linear range of a normalized force-displacement curve.

### Laser ablation

Laser ablation experiments were performed as described previously^54^. Live imaging for laser ablation was performed on an inverted microscope (Eclipse Ti-E; Nikon) with a spinning disk confocal (CSU-X1; Yokogawa Electric Corporation), head dichroic Semrock Di01-T405/488/561GFP, 488 nm (120 mW) and 561 nm (150 mW) diode lasers, emission filters ET525/36M (Chroma Technology Corp.) for GFP or ET630/75M for RFP, and an iXon3 camera (Andor Technology). Targeted laser ablation (20 3-ns pulses at 20 Hz at two target spots) using 551- (for GFP) or 514-nm (for RFP) light was performed using a galvo-controlled MicroPoint Laser System (Photonic Instruments) operated through Metamorph. Laser strength was calibrated before each experimental session based on minimum power necessary to ablate a kinetochore-fiber bundle of microtubules in mitotic cells.

MECs stably expressing LifeAct-RFP-T or inducible ROCK constructs ligated to 2D or 3D ECM were imaged for a short period of time (∼20s) prior to high power ablation of the actin cortex and ∼5 min after laser ablation. The cellular response (bleb nucleation or no response) was quantified. Notably, actomyosin contractility of MECs stably expressing an inducible ROCK construct was induced overnight using 4-hydroxytamoxifen and the actin cortex of these cells was labelled with SiR-actin dye (Cytoskeleton) according to manufacturer’s instructions prior to laser ablation.

### Correlative light and cryo-electron microscopy

No current protocol was available to investigate the role played by matrix stiffness, composition and dimensionality using a combination of correlated light and electron microscopy (CLEM) and quantifiable cryo-electron tomography (cryo-ET) on fully hydrated specimens. To this purpose, we thereby generated a workflow where compliant PA hydrogels of set elastic module were combined with electron microscopy-compatible surfaces functionalized with an ECM component of choice. Details about this novel method are presented in (Gaietta et al manuscript in preparation). Briefly, we prepared two separate modules, one incorporating an electron microscopy compatible surface that could be treated with an ECM component such as Matrigel, and one with a PA hydrogel of set stiffness (i.e. cells cultured under 2D conditions). The two modules were combined right before seeding MCF10A cells expressing mCherry-CAAX (farnesylated mCherry). After a brief incubation, non-adherent cells were removed by gentle aspiration and fresh media containing DMSO (2D control) or blebbistatin (2D and lower cortical tension) was added. Cells were then closely monitored and fixed to halt cellular activity as soon as spreading was noticed. For correlative analysis in CLEM mode, grids were examined on an inverted fluorescent microscope and the position of non-spread cells expressing mcherry-CAAX were recorded. Following the last live-cell image capture, cell movement was halted while maintaining the structural integrity of the site via rapid fixation^55^. Fixed specimens were manually plunge-frozen in liquid nitrogen cooled liquefied ethane to prevent structural collapse or shrinkage associated with dehydration. Vitrified samples then went through a first round of screening on a cryo-TEM to examine the quality of the vitrification process, carbon layer integrity, and the number of cells and regions amenable for cryo-ET. The fluorescence images collected after fixation and before vitrification were aligned with images of the same region obtained by cryo-EM, using a fiducial-less approach we previously published^35^.

### Cellular cryo-tomography and Image reconstruction

Electron cryo-tomography data was taken with a FEI Titan Krios equipped with a Falcon II or Falcon 3CE detector, operated at 300 kV, using Serial EM or the Tomo packages (ThermoFisher Scientific; FEI Company) in batch model^56^. The average dose for a complete tilt series was of about 100-120 e_-_/Å^2^ and the defocus ranged between 8-14 µm. Magnification was chosen to result in pixel sizes of 0.45 nm in the reconstructions. The fidelity and quality of the data collection was monitored with real-time automatic reconstruction protocols implemented in the pyCoAn package, an extended python version of the CoAn package^57^. Briefly, tilt series were aligned using the IMOD package with a combination of fiducial-based and patch-based approaches^58^. Three-dimensional reconstructions were then generated using the Simultaneous Iterative Reconstruction Technique as implemented in Tomo3D^59^. A total of 32 and 28 tomograms near the cell edges were analyzed for cells in the presence and absence of blebbistatin respectively. Protrusion widths were measured in virtual slices of the 3D reconstructions every 100 nm along the center line of the protrusions. Assessment of membrane compartment content was classified into ‘empty’ and ‘filled’ based on the appearance of recognizable macromolecular particles within the compartments. The process was repeated three times to compile statistics. The standard deviation of the assessment was 10.3%. Statistical significance of differences between the distributions was assessed using Mann-Whitney rank tests. The distance between actin filaments in the protrusions was analyzed using a modified version of the sliding-window Fourier amplitude averaging method that adds alignment and classification steps before calculating the Fourier transform^60^.

### Stable isotope labelling with amino acids in cell culture (SILAC), proximity-dependent biotinylation and affinity purification of membrane-associated proteins

MCF10A MECs stably expressing farnesylated APEX2 (APEX2-CAAX) were cultured in lysine- and arginine-free DMEM/F12 media supplemented with dialyzed horse serum (Gemini Bio-Products), EGF, insulin, cholera toxin, hydrocortisone, light isotope labelled lysine and arginine or heavy isotope labelled lysine (K8) and arginine (R10). Cells were labelled in SILAC media for at least 8 passages prior to biotinylation assay. MCF10A were trypsinized and plated on compliant PA gels as described above (2D setup). During cell plating, cells were also preincubated with biotin-phenol overnight due to limited membrane permeability of the chemical in MCF10A cells. Cells were either treated with DMSO or 10 µM blebbistatin for 2 hr prior to the biotinylation reaction. Upon 1 min incubation with H_2_O_2_, cells were washed three times with quenching solution and lysed directly in 2% SDS lysis buffer (2% SDS in 100mM Tris-HCl, pH 8.0). The supernatant was diluted to 0.2% SDS and trichloroacetic acid (TCA) was added to a final concentration of 20% and incubated at 4 °C overnight. After TCA precipitation, protein was pelleted by centrifugation at 13, 000 g for 30 min. Pellets were washed with ice-cold 100% acetone, recentrifuged at 13,000 g for 30 min and air-dried. Pellets were resuspended in 8M guanidine-HCl (GnHCl;Sigma), 100mM Tris-HCl pH 8.0 and incubated at room temperature for 1 hr. Subsequently, samples from light and heavy labelled cells were mixed in equal proportion and the combined samples were diluted to 2.5M GnHCl. Biotinylated proteins were captured on 100 µl of packed high capacity NeutrAvidin sepharose beads (Thermo Fisher) overnight at 4 °C. The pull-down samples were washed five times with large volume of 2.5M Gn-HCl, 100mM Tris-HCl, pH 8.0 prior to mass spectrometry analysis. For immunoblotting, the pull-down samples were washed multiple times in 100mM Tris-HCl followed by washes in RIPA buffer (150 mM NaCl, 1% NP40, 0.5% sodium deoxycholate, 0.1% sodium dodecyl sulphate, 50 mM Tris-HCl, pH 8.0).

### Identification of biotinylated proteins using mass spectrometry

NeutrAvidin sepharose beads and purified proteins were resuspended and mildly denatured in 1M GnHCl, 1mM CaCl_2_, and 100mM Tris pH 8.0, to approximately 200 µL of slurry, where we assumed ∼50% volume occupancy by the beads. Disulfide bonds were reduced with 10 mM tris (2-carboxyethyl) phosphine (Sigma, C4706), and free thiols were alkylated with 40 mM 2-chloroacetamide (Sigma, 22790-250G-F) in 100 mM Tris pH 8.0. Beads were heated to 80 °C for 5 min to denature proteins and then kept at room temperature for 45 min in the dark for protein reduction and alkylation. After alkylation, 5 µg of mass spectrometry (MS) grade trypsin (Fisher, PI90057) dissolved in 5 µL 50 mM acetic acid (Sigma, 45754-100ML-F) was added to proteins on beads, and proteins were digested at room temperature for 20 h, in microcentrifuge tubes rotating on a rotisserie rack. The eluate was transferred to a new tube, acidified to a final concentration of 0.5% trifluoroacetic acid (pH < 3, Sigma, AAA1219822) and desalted by reversed phase C18 solid phase extraction (SPE) cartridge, using a SOLA SPE (Fisher, 03150391). Peptides were eluted from the C18 SPE in 30 µL 50% acetonitrile (ACN, Sigma, 03150391) and 0.1% formic acid (FA, Honeywell, 94318-250ML-F) and then dried in a Genevac EZ-2 on an HPLC setting. Dried peptides were resuspended in 2% acetonitrile, 0.1% formic acid in a bath sonicator for 5 min to a concentration of 0.2 µg/µL before MS analysis.

SILAC-labeled peptides (1 ug) were submitted for nano-LC-MS/MS analysis, using an 83 min reversed phase curved gradient (2.4 – 32% acetonitrile, 0.1% formic acid with a concave curve number 7 in Chromeleon) with a 15 cm Acclaim PepMap 100 C18 analytical column (2 µm beads, 75 µm i.d., Fisher, DX164534), running at 200 nL/min on a Dionex Ultimate 3000 RSLCnano pump, in-line with a hybrid quadrupole-Orbitrap Q-Exactive Plus mass spectrometer (ThermoFisher). The method includes a 13 min segment for the sample to load at 500 nL/min 2.4% ACN, 0.1% FA before the gradient and MS acquisition begins and a 6 minute 80% ACN, 0.08% FA wash step at 500 nL/min after the gradient. For the MS analysis, a data dependent method with a parent ion scan at a resolving power of 70,000 and a top 15 method was used for each replicate, selecting the top 15 most intense peaks for MS/MS using HCD fragmentation (normalized collision energy 27). Dynamic exclusion was activated such that parent ions are excluded from MS/MS fragmentation for 20s after initial selection.

For protein identification and quantification, RAW files were analyzed by Maxquant using default settings^61, 62^. The recorded spectra from two independent biological replicates were searched against the human reference proteome from UniProt (2017-11-15 release, with 71,544 entries in SwissProt/TrEMBL) using MaxQuant, version 1.6.2.1. Search parameters allowed 2 missed tryptic cleavages. Oxidation of methionine, phosphorylation of serine/threonine/tyrosines, and N-terminal acetylation were allowed as variable modifications, while carbamidomethylation of cysteines was selected as a constant modification and a threshold peptide spectrum match (PSM) false discovery rate (FDR) and protein FDR of 1% was allowed. Quantification of SILAC ratios was performed by Maxquant on the MS1 level and the resulting ratios for all replicates were compared using statistical tools found in the Perseus statistical analysis package^63^. Proteins with ratio quantification in only one replicate were removed. Statistical significance was determined by applying a one-sample, two-sided Student T-test to the replicates with a p-value cut off of p = 0.1. The mass spectrometry proteomics data have been deposited to the ProteomeXchange Consortium via the PRIDE partner repository with the dataset identifier PXD018975^64^.

### VSVG trafficking assay

MCF10A cells were transfected with the VSVGts045-GFP construct using Lipofectamine^TM^ 3000 Reagent (Thermo Fisher) for 4hr and subsequently seeded on compliant PA gels. Cells that were unattached after 1hr were removed by media replacement, with fresh media (2D) or media supplemented with rBM (3D). MEC cultures on compliant PA gels were placed at 40 °C overnight to trap the protein at the endoplasmic reticulum. Cells were either fixed immediately or fixed after 2 hr incubation at 32 °C incubation in the presence of cycloheximide. The subcellular localization of VSVG-ts045-GFP was examined using confocal microscopy. Cell surface VSVG and total VSVG fluorescent signals were measured using Fiji software. The trafficking efficiency under different conditions was reported as cell surface fluorescent intensity/total fluorescent signal.

### Immunoblotting

MECs on 2D and 3D PA gels were lysed directly using 2X protein sample buffer (2% sodium dodecyl sulfate, 0.1% bromophenol blue and 20% glycerol, 5% β-mercaptoethanol) containing a protease inhibitor cocktail (Pierce). The lysate was sonicated using multiple short bursts with a stainless steel probe sonicator before boiling at 95 °C for 5 min. Samples were centrifuged at 13 000 g for 10 min and resolved using SDS-PAGE gels and transferred onto Polyvinylidene difluoride membranes (PVDF; BioRad). The membranes were blocked with 5% milk in TBST and probed with the indicated primary antibodies in 5% BSA/TBST and a 1:10000 dilution of anti-rabbit or anti-mouse horseradish peroxidase-conjugated secondary antibodies in 5% BSA/TBST. Blots were developed using WesternBright Quantum (Advansta) and visualized on PXI 6 Touch (Syngene). The band intensity of all western blots was quantified using Fiji software.

### Live/Dead assay

Live/Dead assays of MECs on compliant PA gels were performed using a Viability/Cytotoxicity Kit (Thermo Fisher). Briefly, single round MECs grown in 2D and 3D conditions were seeded for 48hr and were washed twice in PBS prior to live/dead quantification. Live cells were stained using Calcein-AM (Thermo Fisher) and dead cells were stained using Ethidium homodimer-1 (Biotium) according to manufacturer’s instructions. Percent cell death measurements were calculated based on the number of dead cells/total cells. For the 2D monolayer and spheroid live/dead assay, cells were either cultured on 2D plastic surfaces (2D monolayer) or embedded within rBM and grown for 12 days to form 3D spheroids. Cells were treated in the absence or presence of recombinant, purified human or mouse TRAIL (PeproTech), Paclitaxol or doxorubicin. The percent of live/dead cells in 2D monolayers and organoids was determined by quantifying percent active caspase-3 cells by indirect immunofluorescence, normalizing dead cells to total cells enumerated from nuclei counterstained with Hoechst 33342.

### Computational modelling

A summary of the model is described in Supplementary Note 2 and parameters are detailed in Supplementary Table S1.

**Table S1:** Membrane excess area and renormalized surface tension.

| Initial link<br>length | $A/A_P$ | Renormalized $\sigma$ ($k_B T / a_0^2$) | Renormalized $\sigma$<br>($\mu\text{N/m}$) $a_0 = 10$ nm |
| --- | --- | --- | --- |
| 1.27 | 1.0754 | 0.165312855 | 6.794358356 |
| 1.3 | 1.0295 | 0.51881372 | 21.32324391 |
| 1.33 | 1.0156 | 0.85017139 | 34.94204415 |
| 1.35 | 1.0132 | 1.066002153 | 43.81268848 |
| 1.38 | 1.0117 | 1.390551811 | 57.15167943 |
| 1.4 | 1.0112 | 1.619003582 | 66.54104724 |

### Statistical analyses

Statistical analyses were performed using Prism GraphPad 5 software. All quantitative results were assessed by paired or unpaired two-tailed Student’s t-test where indicated or one-way ANOVA after confirming that the data met appropriate assumptions (normality, homogeneous variance, and independent sampling). All data were plotted with standard error bars as indicated in the figure legends, and variances between groups being statistically compared are similar. Sample size was chosen based upon the size of the effect and variance for the different experimental approaches. *P* values less than or equal to 0.05 were considered to be significant (*P<0.05, **P<0.01, ***P< 0.001).

## Supplementary Note 1. Construct source and molecular cloning

The pCB6-VSV-G-ts045-GFP construct and cDNA encoding EGFP-tagged Exo70 were provided by Wei Guo (University of Pennsylvania, USA). The PIP2 reporter (pLEX303-mNeonGreen-PH-PLC) and PIP3 reporter (pLEX303-mNeonGreen-PH-Grp1) constructs were provided by David Bryant (Beatson Institute, UK). The retroviral pBABE-Puro construct with conditionally active human ROCK1 (ROCK1:ER) was provided by Michael Samuel (University of South Australia, Australia). The cDNAs encoding APEX2 and GFP-MAPPER was acquired via Addgene (Plasmid #72480 and #117721, respectively). The cDNAs encoding mApple-golgi (GalT) and human filamin were obtained from Michael Davidson’s plasmid collection at the UCSF Imaging Center (Plasmid #54907 and #54098). pLV-PGK-H2B-EGFP was obtained from Addgene (Plasmid #21210). pCMV-HAhyPBase was kindly provided by Allan Bradley (The Wellcome Trust Sanger Institute, UK). pBMN-LifeAct-RFP-T was provided by Roy Duncan (Dalhousie University, Canada). pLV-PGK-mcherry-CAAX, pLV-rtTAs-M2-IRES-Neo, pPB-SV40-pA-Puro-SV40-polyA-tet-IVS-MCS-BGH-polyA (overexpression vector), and pPB_U6_3xlacIbs_Luciferase_shRNA-IRES_Neo and pPB_U6_3xlacIbs_2kb_stuffer_shRNA-IRES_Neo (shRNA vector) constructs were provided by Johnathon N. Lakins (UCSF, USA). The cDNA encoding IBar-EGFP were provided by Bianxiao Cui (Stanford University, USA). The open reading frame (ORF) of mApple-golgi (GalT) was PCR amplified using forward primer (5’-ACTACTAGATCTACCATGAGGCTTCGGGAGCCGCTC-3’) and reverse primer (5’-TTACTTGTACAGCTCGTCCA-3’) and digested with BglII/BsrGI. The resultant fragment was ligated to a pLV-PGK-H2B-EGFP vector backbone digested with BamHI/BsrGI. EGFP-tagged Exo70 was subcloned into a vector backbone of pLV-PGK-H2B-EGFP digested with BamHI/SalI. A pLKO-cppt-PGK-APEX2-CAAX-IRES-Neomycin construct was generated by ligating three pieces of PCR fragments: (Fragment 1) BamHI-V5-APEX2-N22 (linker)-XhoI; (Fragment 2) XhoI-CAAX-PmeI fragment, and (Fragment 3) pLKO-cppt-hPGK-MCS-IRES-Neomycin construct digested with BamHI/PmeI. The Filamin ORF was PCR amplified using forward primer (5’-ACTACTACTAGTACCATGAGTAGCTCCCACTCTCGGGCG-3’) and reverse primer (5’-ACTACTGTTTAAACTTAGGGCACCACAACGCGGTAGG-3’) and digested with SpeI/PmeI. The resultant fragment was ligated to a pPB-SV40 pA-Puro-SV40-polyA-tet-IVS-vector digested with SpeI/PmeI. For knockdown studies, the shRNAs are cloned into vector pPB_U6_3xlacIbs_2kb_stuffer_shRNA-IRES_Neo digested with AgeI/EcoRI. Filamin shRNA used in this paper has been validated by Sigma Aldrich (TRCN0000230788) and the shRNA is assembled using forward primer: (5’- CCGGGACCGCCAATAACGACAAGAACTCGAGTTCTTGTCGTTATTGGCGGTCTTTTTG-3’) and reverse primer (5’-AATTCAAAAAGACCGCCAATAACGACAAGAACTCGAGTTCTTGTCGTTATTGGCGGTC-3’), which yield duplexes with 5′ AgeI and 3′ EcoRI restriction site overhangs that are ready for ligation reaction. Similarly, Exo70 shRNA is assembled using forward primer: (5’-CCGGCGACCAGCTCACTAAGAACATCTCGAGATGTTCTTAGTGAGCTGGTCGTTTTTTG-3’) and reverse primer 5’-AAT TCA AAA AAC GAC CAG CTC ACT AAG AAC ATC TCG AGA TGT TCT TAG TGA GCT GGT CG-3’).

## Supplementary Note 2. Computational Modelling

### Materials and Methods

The description of modeling methods used for the free energy of tubule formation and free energy landscape for protein recruitment on the cell membranes, as well as the subsequent analysis are detailed in this section.

#### S1.1 Membrane Simulations

Cell membrane simulations consist of a dynamically triangulated sheet evolved with Monte Carlo according to the Helfrich Hamiltonian. This membrane simulation method is adapted from techniques described in Ramakrishnan et al. ^65^. Briefly, in this method the membrane patch is discretized into N vertices, each of characteristic size *a*_0_, interlinked by L links that form T triangles. The membrane is initialized as a planar sheet with periodic boundary conditions, where both the height and curvature of the membrane are maintained over the bounds. Monte Carlo techniques are then used to both evolve membrane morphology and diffuse membrane proteins. The three Monte Carlo moves in the model include the vertex move, which simulates thermal undulations, the link flip, which simulates membrane fluidity and allows it to more dynamically remodel, and the protein move, which allows protein movement along the membrane. All three MC moves are accepted with Metropolis acceptance criteria according to the Helfrich Hamiltonian. The Helfrich Hamiltonian is an energy functional which governs the equilibrium properties of membrane according to,

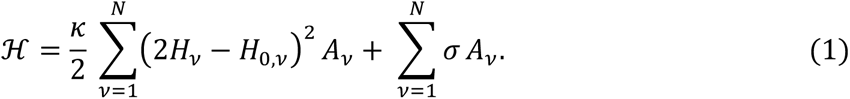

Here *κ* and σ are the bending stiffness and the surface tension of the membrane, *H_v_* and *H_0,v_* are the mean curvature and spontaneous curvatures corresponding to vertex *v,* and *A_v_* is the area associated with vertex *v.* The summation in (1) is computed over all vertices in the discretized membrane. Proteins are included in the model through their spontaneous curvature field *H_0_*. This spontaneous curvature field can be patterned on the membrane to model different biophysical processes. In the case of the generation of cell protrusions, a spontaneous curvature field is assumed to be a circular collection of vertices of a prescribed spontaneous curvature with a nearest-neighbor (Ising-model-like) potential keeping these vertices together. This circular region of spontaneous curvature is meant to approximate the effects of a large complex protein-assembly involved in pushing out a protrusion. As a first approximation, for single membrane proteins, spontaneous curvature fields are assumed to be the form of a Gaussian function with a pre-factor *C*_0_ (peak spontaneous curvature), and a variance *ɛ*^2^ (field spatial extent).

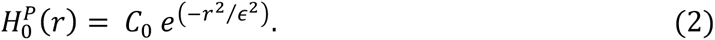

One measure of membrane morphology is the excess area present in the membrane; this quantity is defined simply as the sum of the curvilinear area of the mesh divided by the sum of its projected area on the xy-plane. High excess areas correspond to low tension, while excess areas close to 1 describe tense membranes. For all simulations the bending stiffness was initialized at 20 *k_B_T* and the tension at 0 *k_B_T/*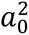, where *T* is the temperature and *a_0_* sets the link-length between adjacent vertices. Both these moduli are renormalized during equilibration for different initial conditions of link length. Renormalized tension is calculated by fitting the undulation spectrum of membrane undulations ^33^. A table of the renormalized values of tension is shown in Table S1. A typical value of a_0_ for the systems we model is a_0_=10 nm; this value ensures that our choices of c_0_=0.4-0.8 a_0_^-1^ and ε^2^=6.3a_0_^2^, are primed to model curvature sensing proteins such with BAR and ENTH domains, as justified in previous works ^32, 33^. With this choice of the value for the scale parameter a_0_, the characteristic size of the protrusion we model is of diameter 10a_0_ (as set by the Ising potential), which amounts to 100 nm; this value is consistent with the diameter of membrane protrusions observed in cell studies [W. Guo, private communication, 2015]. We note that the choice of a_0_ also sets the range of membrane tensions we model to the range 0-100 µN/m, which is consistent with the range to membrane tension values reported in cellular experiments ^66, 67^.

#### S1.2 Free Energy Methods

The membrane model is coupled to several computational free energy methods including Widom insertion, inhomogeneous Widom insertion, and thermodynamic integration ^32, 33, 65^.

We employ Widom insertion as a computational method used to probe the excess chemical potential of a curvature inducing protein on a membrane with a given tension/excess area. The Widom insertion involves randomly inserting a virtual/test membrane protein into the system and recording the change in energy. This difference in energy between a system with n and n+1 proteins can be related to the excess chemical potential as,

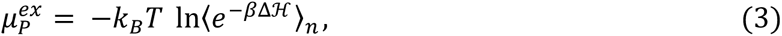

where 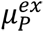 is the excess chemical potential for the n+1’th protein and 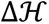 is the change in energy calculated from the Helfrich Hamiltonian upon insertion of that protein. The Boltzmann weighted ensemble average of this change in energy is then related to the chemical potential. Extensive sampling of each equilibrated system can then converge this ensemble average. In all Widom results shown, simulations were equilibrated for 5 million Monte Carlo steps and sampled with Widom method every 100 steps. Only the last 2.5 million Monte Carlo steps are used when computing the ensemble average to ensure an equilibrated system for sampling.

Widom insertion can also be derived in a spatially dependent manner where subsections of the system can be isolated by an axis of inhomogeneity. In inhomogeneous Widom insertion, each subsection has its excess chemical potential defined as

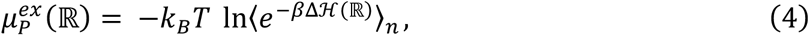

where ℝ is a collection of vertexes within a specified region. In the case of protrusion simulations, where a circular region induces spontaneous curvature, three regions are defined: a protrusion region, defined as all vertices within the Ising-potential, an annulus region, defined as a radial extension of the protrusion region, and a basal region consisting of the rest of the vertices in the membrane, see Figure S1.

Thermodynamic Integration (TI) is a computational free energy perturbation method that computes the change in free energy between two states of the system (states A and B). TI works by defining a scalar parameter λ, which traces a path between these two states (state A: λ = 1; state B: λ = 0). The derivative of the Helmholtz free energy *F = -k_B_T* In *Q* (*Q* is the partition function) with respect to the parameter λ is given by:

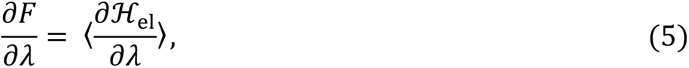

and the free energy difference between the states characterized by *λ* = 0 and *λ* = 1 is calculated as,

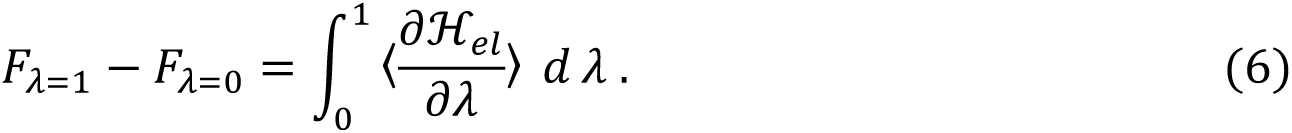

The change in free energy is calculated in the TI method by running a set of simulations with a range of lambda values from 0 to 1, where the integrand in equation (6) is calculated and stored. The discretized data are then integrated with the trapezoid rule to obtain the free energy difference between *λ* = 0 and *λ* = 1. All results shown were obtained with a window resolution of 0.1 (or *λ* = 0, 0.1, 0.2, … ., 1). In order to determine the accuracy of this window resolution, a sensitivity analysis was performed, where window sizes of 0.05 and 0.02 showed no appreciable difference in free energy.

Error was estimated in all free energy methods by running each simulation in quadruplicate and calculating the standard deviation.

#### S1.3 Reference State of Free Energy of Membrane Protrusions

To generate membrane protrusions, a spontaneous curvature field is applied at the center of the membrane that consists of 50 vertices; this gives the protrusion region an average radius of 4.55 *a*_0_. The spontaneous curvature of the protrusion was chosen to be either 0.4 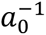 or 0.6 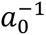 these values of spontaneous curvature gave the largest variety of protrusion structures when altering the membrane tension. Simulations with the spontaneous curvature 0.6 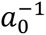 and low tension showed the generation of budding vesicles, while all other simulations generated protrusion-like morphologies without a constricted vesicle neck.

The free energy of formation of a protrusion was analyzed with TI. As described above, this method computes the change in free energy to deform the membrane with an applied spontaneous curvature field. In order to analyze the free energy of generating a protrusion on the membrane, the reference state of the free energy is changed according to,

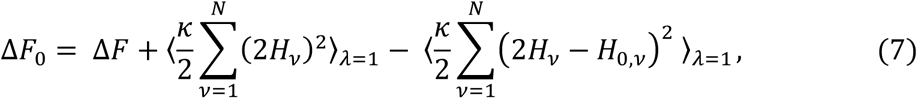

where Δ*F* is the change in free energy required to deform the membrane once the protrusion is assembly is already bound, and Δ*F*_0_ is the change in free energy to generate a protrusion from an unstressed membrane.

### S2. Studies of Protein Recruitment on Protrusions

Membrane protein recruitment on protrusions was analyzed by using inhomogeneous Widom insertion techniques coupled with simulation conditions detailed in the previous section (S1.3). In this study membrane protrusion morphologies are generated by applying a spontaneous curvature field of 0.4 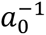 in a circular pattern with an average radius of 4.55 *a*_0_, as in the previous section. This study is conducted with no membrane proteins present (i.e. n = 0, n+1 = 1). Inhomogeneous Widom insertion is then done in three spatially distinct regions: protrusion, annulus, and basal regions, see Figure S1. A snapshot of the simulations with panels depicting the three membrane regions, and a height map of the protrusion is shown in Figure S1.

When conducting inhomogeneous Widom calculations, the background spontaneous curvature field of the protrusion-field was disregarded in the Widom formula. Membrane proteins with spontaneous curvature fields with the functional form in equation (2) were then randomly inserted and categorized into the appropriate region. The protrusion region was defined as all 50 vertices with the spontaneous curvature field. The annulus region was defined by creating a neighbor list at the point of insertion: if a protrusion region vertex existed within the first 30 closest neighbors of this vertex and the vertex was not a protrusion vertex itself, this was described as the annulus region vertex. Vertices that don’t meet either of these criterions were categorized as the basal region. The proteins inserted in this study had peak spontaneous curvature of *C*_0_ = 0.4 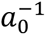 or −0.4 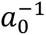 and a variance of *ɛ*^2^ = 6.3 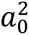.

### S3. Protrusion Elongation Studies

Protrusion simulations are performed with several curvature-inducing membrane proteins present and with no other background spontaneous curvature field. In this study, membrane proteins diffuse around and are able to co-locate to produce tubule like structures. Once above a threshold density required for tubulation, the protrusion elongates. The threshold density of proteins required for tubulation and consequently the protrusion length and density are strongly dependent on membrane interfacial tension.

---

**Figure S1:**
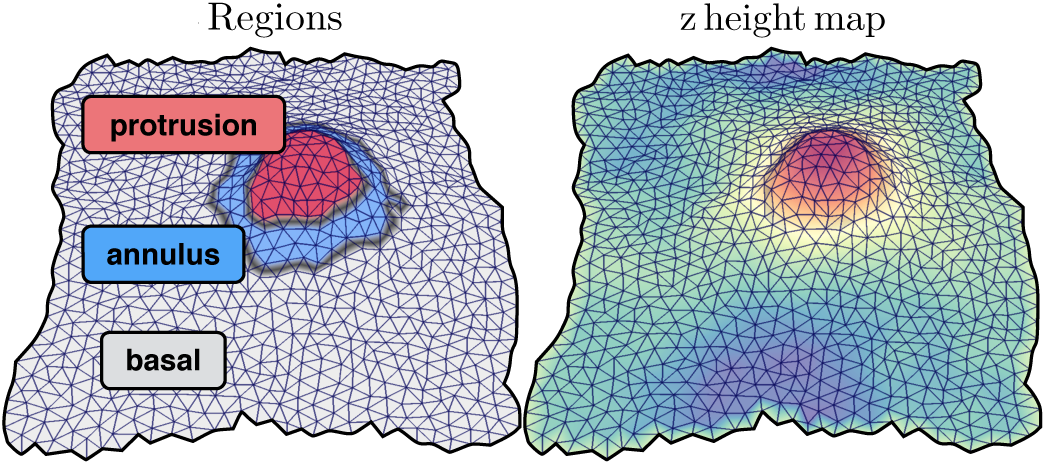
Snapshot of membrane simulation detailing (left) the three inhomogeneous Widom regions, and (right) a heightmap of the z-axis of a membrane protrusion.

**Figure S4.**
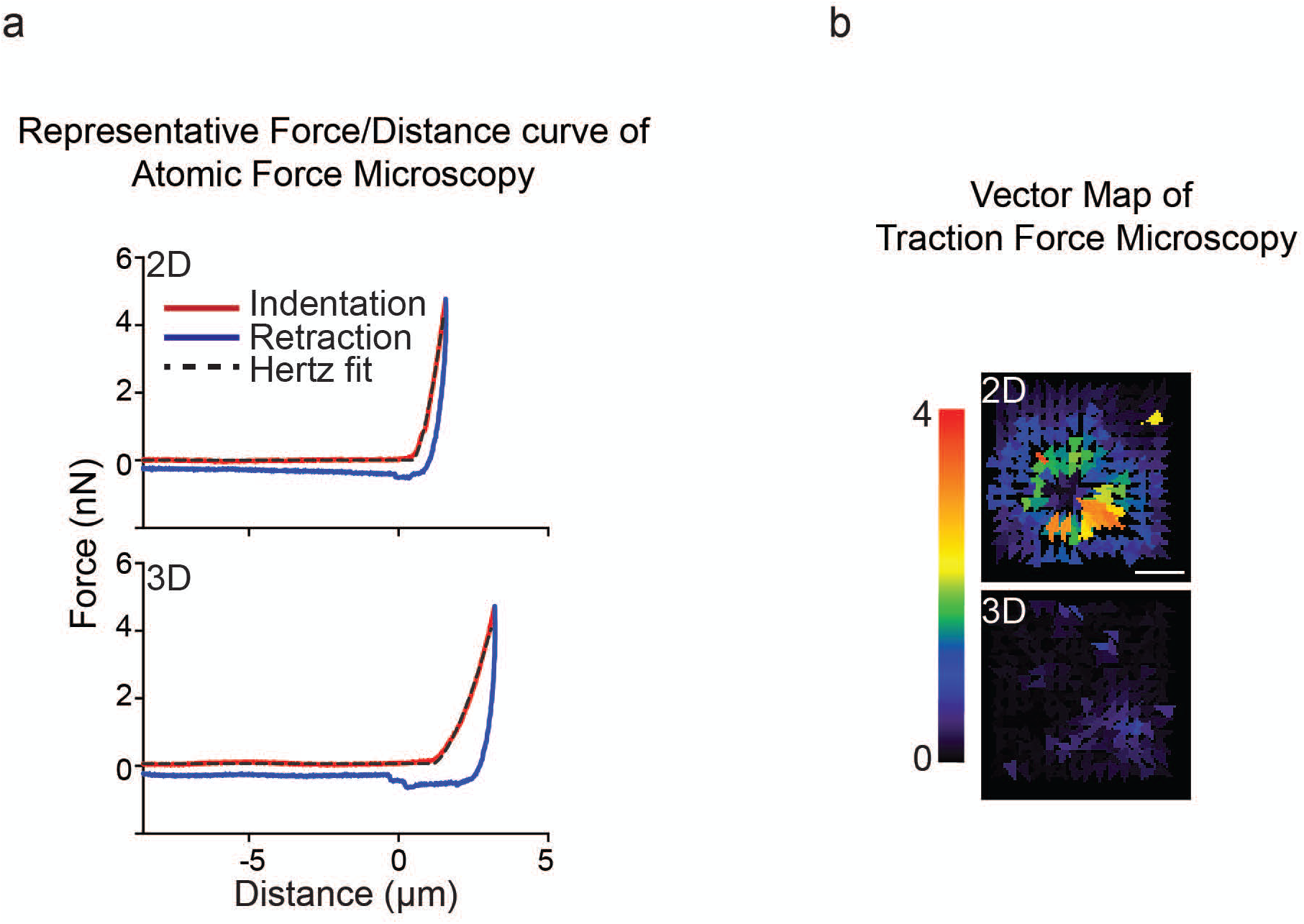
rBM ligation in 3D reduces cellular cortical actin tension. **(a)** Bead displacement field of MECs ligated to a rBM in 2D or 3D is displayed as a color-coded vector map. Scale bar, 10 μm. **(b)** Representative AFM force-distance curve of MECs interacting with laminin-111 in either 2D or 3D. The indentation curve (red), retraction curve (blue), and Hertz model fit of the indentation curve (black dash line) are shown.

**Figure S6.**
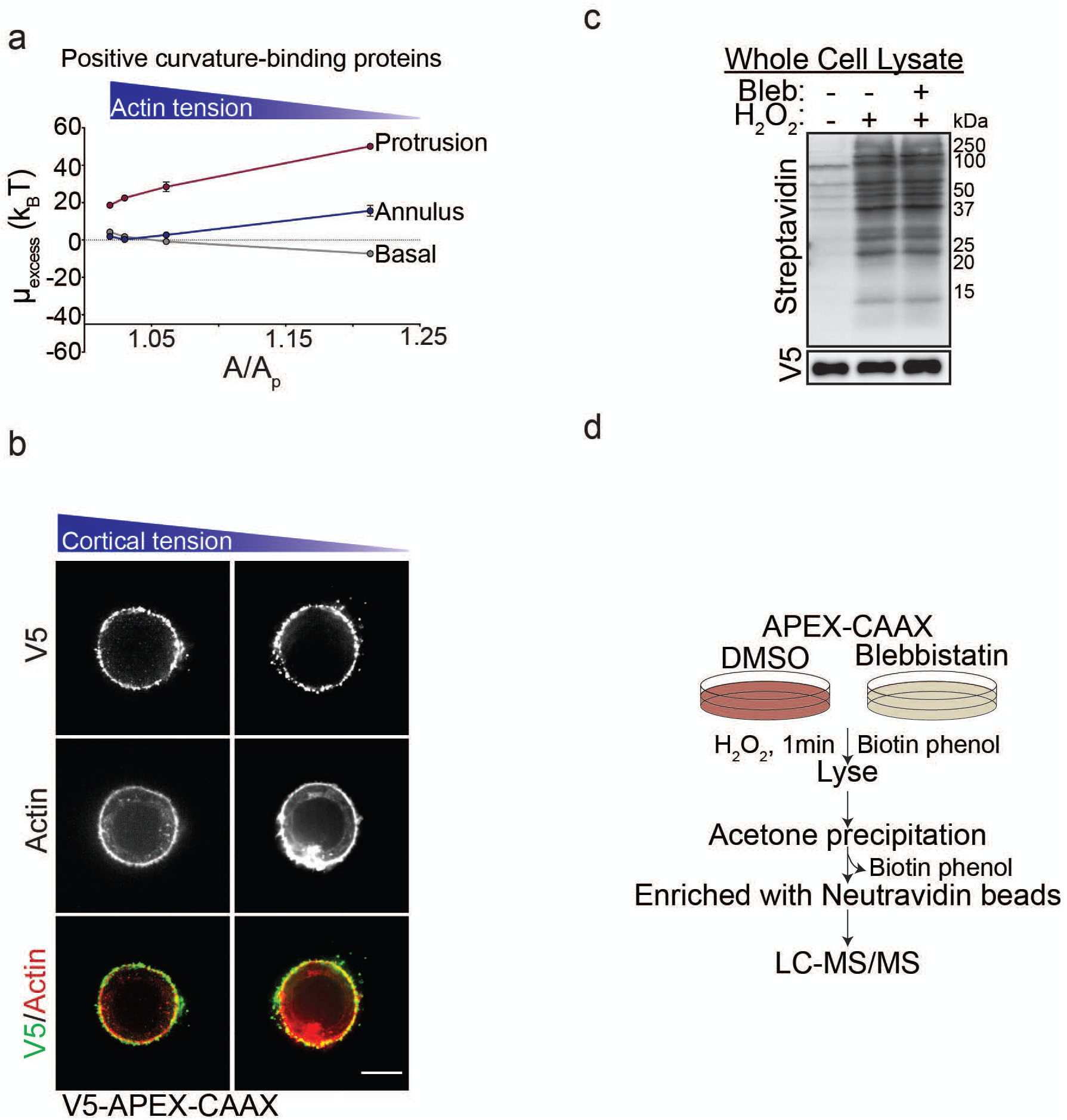
Cortical actin tension modulates plasma membrane protein composition. **(a)** Excess chemical potential required for the recruitment of positive curvature sensing domains in basal, annulus, and protrusion regions of the plasma membrane plotted as a function of A/Ap. **(b)** Representative fluorescence microscopy images of MECs stably expressing recombinant V5-APEX2-CAAX and ligated to rBM in 2D with and without blebbistatin treatment (to reduce cortical tension). MECs were immunostained with antibodies targeting V5 (Alexa488; green) and counterstained with Texas-red phalloidin (red). Scale bar, 10 μm. **(c)** MECs ligated to rBM in 2D with and without 2 hr of blebbistatin treatment were harvested for immunoblotting. The biotinylated proteins were detected using streptavidin-HRP. **(d)** Cartoon depicting strategy used for the two-state SILAC experiment. MECs expressing APEX-CAAX were treated with biotin-phenol overnight followed by 1 min of H_2_O_2_ exposure. MECs labelled with heavy isotope amino acids were treated with blebbistatin to reduce myosin II activity whereas those labelled with light amino acids were treated with DMSO (vehicle). Cells were lysed and excess biotin phenol trapped in the polyacrylamide gels was removed using acetone precipitation. The resuspended protein was purified using streptavidin beads and identified by mass spectrometry. For each protein, the H/L SILAC ratio reflects the extent of its biotinylation by APEX2-CAAX in the presence/absence of blebbistatin.

## References

1. Roignot, J., Peng, X. & Mostov, K. Polarity in mammalian epithelial morphogenesis. Cold Spring Harb Perspect Biol 5 (2013).

2. Weaver, V.M. et al. beta4 integrin-dependent formation of polarized three-dimensional architecture confers resistance to apoptosis in normal and malignant mammary epithelium. Cancer Cell 2, 205–216 (2002).

3. Han, K. et al. CRISPR screens in cancer spheroids identify 3D growth-specific vulnerabilities. Nature 580, 136–141 (2020).

4. Chen, L.H. & Bissell, M.J. A novel regulatory mechanism for whey acidic protein gene expression. Cell Regul 1, 45–54 (1989).

5. Streuli, C.H., Bailey, N. & Bissell, M.J. Control of mammary epithelial differentiation: basement membrane induces tissue-specific gene expression in the absence of cell-cell interaction and morphological polarity. J Cell Biol 115, 1383–1395 (1991).

6. Roskelley, C.D., Desprez, P.Y. & Bissell, M.J. Extracellular matrix-dependent tissue-specific gene expression in mammary epithelial cells requires both physical and biochemical signal transduction. Proc Natl Acad Sci U S A 91, 12378–12382 (1994).

7. Paszek, M.J. et al. Tensional homeostasis and the malignant phenotype. Cancer Cell 8, 241–254 (2005).

8. Krebs, J., Agellon, L.B. & Michalak, M. Ca(2+) homeostasis and endoplasmic reticulum (ER) stress: An integrated view of calcium signaling. Biochem Biophys Res Commun 460, 114–121 (2015).

9. Lewis, R.S. Store-operated calcium channels: new perspectives on mechanism and function. Cold Spring Harb Perspect Biol 3 (2011).

10. Petersen, O.W., Ronnov-Jessen, L., Howlett, A.R. & Bissell, M.J. Interaction with basement membrane serves to rapidly distinguish growth and differentiation pattern of normal and malignant human breast epithelial cells. Proc Natl Acad Sci U S A 89, 9064–9068 (1992).

11. Streuli, C.H. & Bissell, M.J. Expression of extracellular matrix components is regulated by substratum. J Cell Biol 110, 1405–1415 (1990).

12. Barcellos-Hoff, M.H., Aggeler, J., Ram, T.G. & Bissell, M.J. Functional differentiation and alveolar morphogenesis of primary mammary cultures on reconstituted basement membrane. Development 105, 223–235 (1989).

13. van Vliet, A.R. et al. The ER Stress Sensor PERK Coordinates ER-Plasma Membrane Contact Site Formation through Interaction with Filamin-A and F-Actin Remodeling. Mol Cell 65, 885–899 e886 (2017).

14. Razinia, Z., Makela, T., Ylanne, J. & Calderwood, D.A. Filamins in mechanosensing and signaling. Annu Rev Biophys 41, 227–246 (2012).

15. Guo, M. et al. Cell volume change through water efflux impacts cell stiffness and stem cell fate. Proc Natl Acad Sci U S A 114, E8618–E8627 (2017).

16. Han, Y.L. et al. Cell swelling, softening and invasion in a three-dimensional breast cancer model. Nature Physics 16, 101–108 (2020).

17. Tinevez, J.Y. et al. Role of cortical tension in bleb growth. Proc Natl Acad Sci U S A 106, 18581–18586 (2009).

18. Kitamura, M. & Hiramatsu, N. Real-time monitoring of ER stress in living cells and animals using ESTRAP assay. Methods Enzymol 490, 93–106 (2011).

19. Bisaria, A., Hayer, A., Garbett, D., Cohen, D. & Meyer, T. Membrane-proximal F-actin restricts local membrane protrusions and directs cell migration. Science 368, 1205–1210 (2020).

20. Jarsch, I.K., Daste, F. & Gallop, J.L. Membrane curvature in cell biology: An integration of molecular mechanisms. J Cell Biol 214, 375–387 (2016).

21. Ramakrishnan, N., Sunil Kumar, P.B. & Radhakrishnan, R. Mesoscale computational studies of membrane bilayer remodeling by curvature-inducing proteins. Phys Rep 543, 1–60 (2014).

22. Tourdot, R.W., Bradley, R.P., Ramakrishnan, N. & Radhakrishnan, R. Multiscale computational models in physical systems biology of intracellular trafficking. IET Syst Biol 8, 198–213 (2014).

23. Tourdot, R.W., Ramakrishnan, N. & Radhakrishnan, R. Defining the free-energy landscape of curvature-inducing proteins on membrane bilayers. Phys Rev E Stat Nonlin Soft Matter Phys 90, 022717 (2014).

24. Volkmann, N., DeRosier, D., Matsudaira, P. & Hanein, D. An atomic model of actin filaments cross-linked by fimbrin and its implications for bundle assembly and function. J Cell Biol 153, 947–956 (2001).

25. Anderson, K.L. et al. Nano-scale actin-network characterization of fibroblast cells lacking functional Arp2/3 complex. J Struct Biol 197, 312–321 (2017).

26. McMahon, H.T. & Boucrot, E. Membrane curvature at a glance. J Cell Sci 128, 1065–1070 (2015).

27. Zhao, Y. et al. Exo70 generates membrane curvature for morphogenesis and cell migration. Dev Cell 26, 266–278 (2013).

28. Umbrecht-Jenck, E. et al. S100A10-mediated translocation of annexin-A2 to SNARE proteins in adrenergic chromaffin cells undergoing exocytosis. Traffic 11, 958–971 (2010).

29. Boye, T.L. et al. Annexins induce curvature on free-edge membranes displaying distinct morphologies. Sci Rep 8, 10309 (2018).

30. Walter, P. & Ron, D. The unfolded protein response: from stress pathway to homeostatic regulation. Science 334, 1081–1086 (2011).

31. Zhu, Y., Wu, B. & Guo, W. The role of Exo70 in exocytosis and beyond. Small GTPases 10, 331–335 (2019).

32. Kim, T.J. et al. Distinct mechanisms regulating mechanical force-induced Ca(2)(+) signals at the plasma membrane and the ER in human MSCs. Elife 4, e04876 (2015).

33. Lee, N.S. et al. Focused Ultrasound Stimulates ER Localized Mechanosensitive PANNEXIN-1 to Mediate Intracellular Calcium Release in Invasive Cancer Cells. Front Cell Dev Biol 8, 504 (2020).

34. Nava, M.M. et al. Heterochromatin-Driven Nuclear Softening Protects the Genome against Mechanical Stress-Induced Damage. Cell 181, 800–817 e822 (2020).

35. Maiers, J.L. & Malhi, H. Endoplasmic Reticulum Stress in Metabolic Liver Diseases and Hepatic Fibrosis. Semin Liver Dis 39, 235–248 (2019).

36. Sequeira, S.J. et al. Inhibition of proliferation by PERK regulates mammary acinar morphogenesis and tumor formation. PLoS One 2, e615 (2007).

## References

50. Weaver, V.M. et al. Reversion of the malignant phenotype of human breast cells in three-dimensional culture and in vivo by integrin blocking antibodies. J Cell Biol 137, 231–245 (1997).

51. Przybyla, L., Lakins, J.N., Sunyer, R., Trepat, X. & Weaver, V.M. Monitoring developmental force distributions in reconstituted embryonic epithelia. Methods 94, 101–113 (2016).

52. Masters, T. et al. Easy fabrication of thin membranes with through holes. Application to protein patterning. PLoS One 7, e44261 (2012).

53. Hutter, J.L. & Bechhoefer, J. Calibration of atomic-force microscope tips. Review of Scientific Instruments 64, 1868–1873 (1993).

54. Long, A.F., Udy, D.B. & Dumont, S. Hec1 Tail Phosphorylation Differentially Regulates Mammalian Kinetochore Coupling to Polymerizing and Depolymerizing Microtubules. Curr Biol 27, 1692–1699 e1693 (2017).

55. Marston, D.J. et al. High Rac1 activity is functionally translated into cytosolic structures with unique nanoscale cytoskeletal architecture. Proc Natl Acad Sci U S A 116, 1267–1272 (2019).

56. Mastronarde, D.N. Automated electron microscope tomography using robust prediction of specimen movements. J Struct Biol 152, 36–51 (2005).

57. Volkmann, N. & Hanein, D. Quantitative fitting of atomic models into observed densities derived by electron microscopy. J Struct Biol 125, 176–184 (1999).

58. Mastronarde, D.N. & Held, S.R. Automated tilt series alignment and tomographic reconstruction in IMOD. J Struct Biol 197, 102–113 (2017).

59. Agulleiro, J.I. & Fernandez, J.J. Fast tomographic reconstruction on multicore computers. Bioinformatics 27, 582–583 (2011).

60. Kumar, A. et al. Local Tension on Talin in Focal Adhesions Correlates with F-Actin Alignment at the Nanometer Scale. Biophys J 115, 1569–1579 (2018).

61. Cox, J. & Mann, M. MaxQuant enables high peptide identification rates, individualized p.p.b.-range mass accuracies and proteome-wide protein quantification. Nat Biotechnol 26, 1367–1372 (2008).

62. Tyanova, S., Temu, T. & Cox, J. The MaxQuant computational platform for mass spectrometry-based shotgun proteomics. Nat Protoc 11, 2301–2319 (2016).

63. Tyanova, S. et al. The Perseus computational platform for comprehensive analysis of (prote)omics data. Nat Methods 13, 731–740 (2016).

64. Perez-Riverol, Y. et al. The PRIDE database and related tools and resources in 2019: improving support for quantification data. Nucleic Acids Res 47, D442–D450 (2019).

65. Ramakrishnan, N., Sunil Kumar, P.B. & Radhakrishnan, R. Mesoscale computational studies of membrane bilayer remodeling by curvature-inducing proteins. Physics Reports 543, 1–60 (2014).

66. Diz-Muñoz, A., Fletcher, D.A. & Weiner, O.D. Use the force: membrane tension as an organizer of cell shape and motility. Trends in Cell Biology 23, 47–53 (2013).

67. Vogel, V. & Sheetz, M. Local force and geometry sensing regulate cell functions. Nat Rev Mol Cell Biol 7, 265–275 (2006).

